# KSHV promotes oncogenic FOS to inhibit nuclease AEN and transactivate RGS2 for AKT phosphorylation

**DOI:** 10.1101/2024.01.27.577582

**Authors:** Vladimir Majerciak, Beatriz Alvarado-Hernandez, Yanping Ma, Shivalee Duduskar, Alexei Lobanov, Maggie Cam, Zhi-Ming Zheng

## Abstract

Kaposi’s sarcoma-associated herpesvirus (KSHV) ORF57 is a lytic RNA-binding protein. We applied BCBL-1 cells in lytic KSHV infection and performed UV cross-linking immunoprecipitation (CLIP) followed by RNA-seq of the CLIPed RNA fragments (CLIP-seq). We identified ORF57-bound transcripts from 544 host protein-coding genes. By comparing with the RNA-seq profiles from BCBL-1 cells with latent and lytic KSHV infection and from HEK293T cells with and without ORF57 expression, we identified FOS and CITED2 RNAs being two common ORF57-specific RNA targets. FOS dimerizes with JUN as a transcription factor AP-1 involved in cell proliferation, differentiation, and transformation. Knockout of the ORF57 gene from the KSHV genome led BAC16-iSLK cells incapable of FOS expression in KSHV lytic infection. The dysfunctional KSHV genome in FOS expression could be rescued by Lenti-ORF57 virus infection. ORF57 protein does not regulate FOS translation but binds to the 13-nt RNA motif near the FOS RNA 5ʹ end and prolongs FOS mRNA half-life 7.7 times longer than it is in the absence of ORF57. This binding of ORF57 to FOS RNA is competitive to the binding of a host nuclease AEN (also referred to as ISG20L1). KSHV infection inhibits the expression of AEN, but not exosomal RNA helicase MTR4. FOS expression mediated by ORF57 inhibits *AEN* transcription, but transactivates *RGS2,* a regulator of G-protein coupled receptors. FOS binds a conserved AP-1 site in the *RGS2* promoter and enhances RGS2 expression to phosphorylate AKT. Altogether, we have discovered that KSHV ORF57 specifically binds and stabilizes FOS RNA to increase FOS expression, thereby disturbing host gene expression and inducing pathogenesis during KSHV lytic infection.

**Significance:** We discovered that FOS, a heterodimer component of oncogenic transcription factor AP- 1, is highly elevated in KSHV infected cells by expression of a viral lytic RNA-binding protein, ORF57, which binds a 13-nt RNA motif near the FOS RNA 5ʹ end to prolong FOS RNA half-life. This binding of ORF57 to FOS RNA is competitive to the binding of host RNA destabilizer(s). KSHV infection inhibits expression of host nuclease AEN (or ISG20L1), but not MTR4. FOS inhibits *AEN* transcription, but transactivates *RGS2* by binding to a conserved AP-1 site in the *RGS2* promoter, thereby enhancing RGS2 expression and phosphorylation of AKT. Our data conclude that viral RNA-binding protein ORF57 controls the expression of a subset of genes for signaling, cell cycle progression, and proliferation to contribute viral pathogenesis.

## INTRODUCTION

Kaposi’s sarcoma-associated herpesvirus (KSHV) is a γ-herpesvirus commonly linked to three types of cancer: Kaposi’s sarcoma, body cavity-based lymphoma, and multicentric Castleman’s disease (1-3). KSHV infection consists of two life cycles: latent and lytic. Expression of viral latent and lytic genes contributes KSHV pathogenesis and tumorigenesis (4).

The well-known proto-oncoprotein FOS (also known as c-FOS) is a nuclear phosphoprotein. FOS homodimerizes with FOS or FOSB or heterodimerizes with JUN or JUNB to form a notorious transcriptional activator protein 1 or AP-1 (5-7). AP-1 expression is rapidly and transiently induced in different cell types by diverse stimuli, including growth factors and bacterial or viral infections (8). In turn, AP-1 regulates cellular processes such as cell proliferation, differentiation, apoptosis, and tumorigenesis. AP-1 also regulates the gene expression of oncogenic and non-oncogenic viruses, including KSHV, EBV, cytomegalovirus, and herpes simplex viruses (9). During KSHV lytic infection, FOS induces the expression of several KSHV proteins, including RTA, a master transactivator of KSHV lytic replication, and K8, a regulator of KSHV lytic DNA replication, by the direct binding to their promoters (10).

FOS expression is regulated at transcriptional, posttranscriptional, and post-translational levels (11). Previous studies suggested the AP-1 or FOS/JUN upregulation during KSHV lytic infection by RTA activation (10) and the increased AP-1 phosphorylation (10) by the MAPK pathway (12-14) or the sustained ERK-RSK activation by KSHV ORF45 (15). AP-1 binds to viral promoters of RTA, K8, ORF57, and IL6 and increases the transcription of both viral and host genes (10, 12, 13). However, no viral protein has been found to directly regulate AP1 expression post-transcriptionally.

The viral RNA-binding protein ORF57 (mRNA transcript accumulation or MTA) is encoded by a viral early lytic gene and is essential for the expression of KSHV lytic genes and productive lytic replication (16-25). ORF57 is a multifunctional homodimeric regulator of viral RNA stability, alternative RNA splicing, and translation, and of host antiviral RNA granules, through direct interactions with viral RNA targets and host cofactors (16, 22, 25-42) and regulates the expression of viral and host coding and noncoding genes (43- 46). Through interaction with hTREX, interestingly, ORF57 in overexpression was also observed to induce the formation of RNA:DNA hybrids and DNA damage (47). Moreover, an early study suggested that viral ORF57 might also regulate transcription in a direct or indirect mechanism (48).

In this report, we provide further evidence of ORF57 direct regulation of the expression of host protein-coding *FOS* gene and contributions to increased expression of FOS RNA and protein during KSHV lytic infection by two mechanisms: First, ORF57 protein binds to a 5ʹ end motif of FOS RNA to protect FOS RNA from rapid decay and enhance FOS expression. Secondly, ORF57-mediated expression of FOS inhibits transcription of *AEN* [apoptosis enhancing nuclease or ISG20L1 (49, 50)], a mediator of p53 in apoptosis induction (51) and a destabilizer of FOS RNA. In contrast, ORF57- mediated FOS upregulation led to direct transcriptional activation of a multifunctional regulator of G-protein signaling 2 (*RGS2*) (52, 53) and phosphorylation of AKT in KSHV-infected cells. Thus, ORF57 transactivates gene transcription indirectly through FOS, a major heterodimer component of transcription factor AP-1.

## RESULTS

### ORF57 associates with host protein-coding RNAs during KSHV lytic infection

KSHV ORF57 protein is highly expressed during KSHV lytic replication. To identify host RNAs associated with ORF57 in KSHV lytic infection for their post-transcriptional regulation, we generated a series of anti-ORF57 polyclonal and monoclonal antibodies against an ORF57 N-terminal epitope (aa 119-PEKRPRRRPRDRLQ-132) and a C-terminal epitope (aa 394-ARGQELFRTLLEYYRPGDV-412) in collaboration with two NCI partners, Rockland Immunochemicals for rabbit polyclonal antibody production and BD Biosciences for mouse monoclonal antibody production. The affinity-purified anti-ORF57 N-terminal epitope antibody (Fig. 1A, left panel) was found highly specific towards ORF57 in Western blot when comparing to the affinity-purified anti-ORF57 C-terminal peptide antibody (Fig. 1A, right panel). Two additional mouse monoclonal antibodies against the ORF57 N-terminal epitope (L36-724.86.254 and L36-1121.207.6) were reported in our previous publications (29, 38) and commercialized by two separate companies. Subsequently, the affinity-purified rabbit polyclonal anti-ORF57 N-terminal antibody was chosen for ORF57 CLIP-seq (cross-linking immunoprecipitation combined by RNA-sequencing) to enrich covalently cross-linked ORF57-RNA complexes from UV-irradiated BCBL-1 cells with lytic KSHV infection. The efficiency of CLIPed ORF57-RNA complexes were confirmed by detection of ORF57 protein by Western blot (Fig. 1B), whereas the ORF57-bound and protected RNA fragments from RNase digestion were extracted for RNA-seq analysis as described in detail in our previous publications (46, 54) (Fig. 1C).

**Figure 1.**
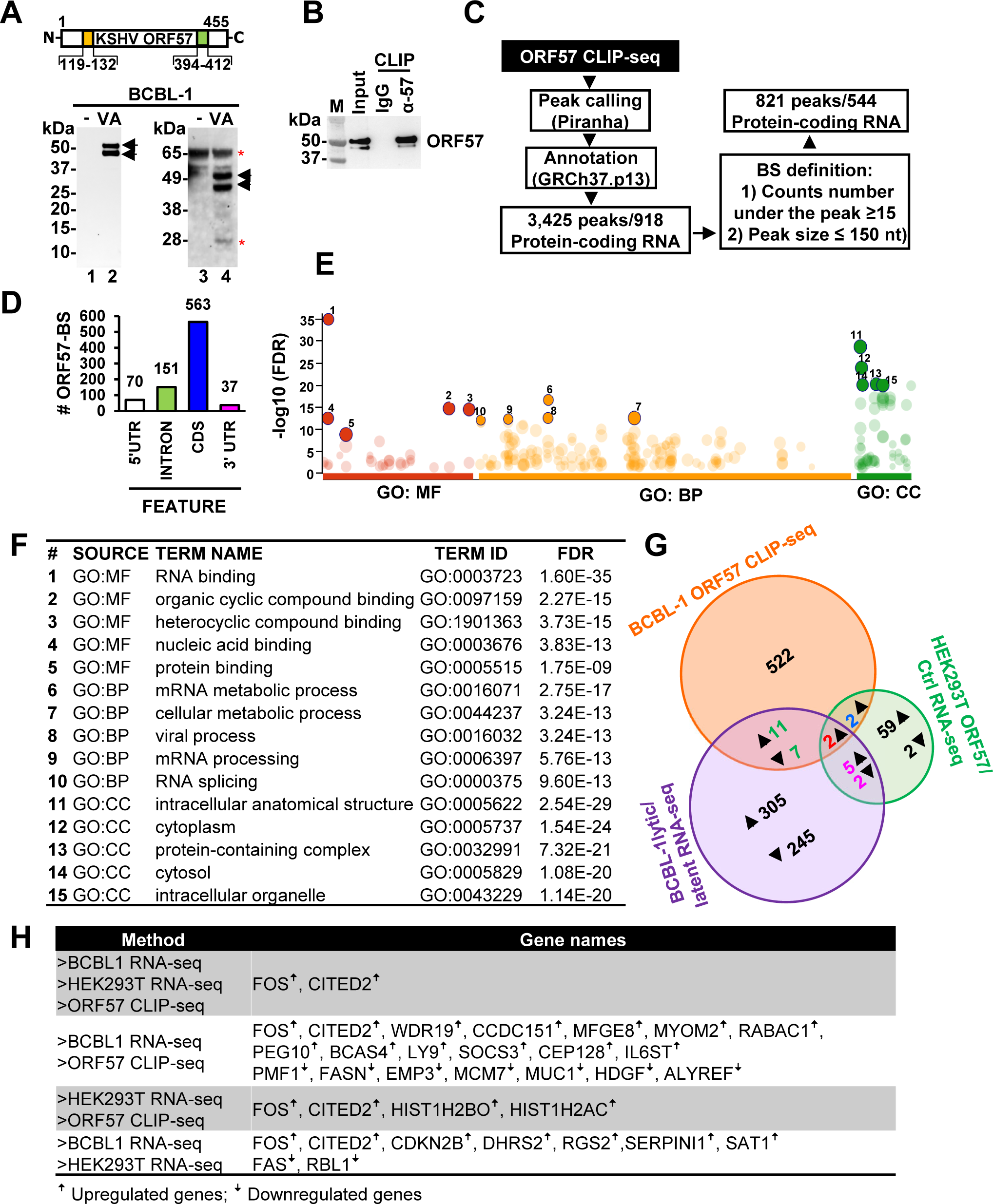
ORF57 preferentially binds to coding sequences in protein-coding RNAs. (A) A diagram showing positions of the epitopes in the ORF57 N-terminus (yellow box) and C-terminus (green box) used for the development of ORF57-specific antibodies. ORF57 detection from BCBL-1 cells without (-) valproic acid (VA) and with (+) VA treatment to induce KSHV lytic infection by Western blot using an affinity-purified ORF57 antibody against the N-terminal (lines 1 and 2) or C-terminal (lines 3 and 4) epitopes. (B and C) ORF57 CLIP-seq using the affinity-purified anti-ORF57 antibody against the N-terminal peptide and subsequent downstream bioinformatics analyses. ORF57 detection by Western blot of the CLIPed ORF57-RNA complexes using the anti-ORF57 antibody along with nonspecific IgG serving as a negative control (B). (C) Bioinformatics workflow in identification of the genome-wide ORF57 binding sites (BS) in RNA transcripts of 544 host protein-coding genes. (D) Number of ORF57 BS mapped to the RNA 5ʹ untranslated region (5ʹ UTR), intron, coding sequences (CDS), and 3ʹ untranslated region (3ʹ UTR). (E and F) Enrichment analysis using the Ensembl ID from 544 host protein-coding gene transcripts containing ORF57 BS. Dot plot of the genes enriched by Gene Ontology (GO) terms for molecular function (MF), biological process (BP), and cellular component (CC). Numbered red, yellow, and dark green dots are the top 5 most enriched GO terms for MF, BP and CC based on the lowest false discovery rate (FDR) in the ranking order of 1-15 (E), which are detailed in (F). (G) Venn diagrams depicting the overlap of highly expressed (RPKM ≥3) protein-coding genes with significantly altered expression change (at least two-fold, p<0.001) identified by RNA-seq of BCBL-1 cells with KSHV lytic versus latent infection, and HEK293T cells with ORF57 versus without ORF57 expression, and host protein-coding transcripts with ORF57 BS identified by ORF57 CLIP-seq. Up and down arrows indicate upregulation or downregulation and the associated numbers of the affected genes. (H) Differentially expressed genes overlapped between individual datasets from the Venn diagrams (G).

By mapping all high-quality RNA-seq reads obtained from three independent anti-ORF57 CLIP-seq experiments (NCBI GEO accession number: <u>GSE179726</u>) to the chimeric human genome GRCh37-KSHV genome (46) and peak calling by Piranha software (version 4.0.164) (55)(https://github.com/smithlabcode/piranha), we identified 1857 ORF57-binding sites (BS) from 1271 human coding and non-coding genes, of which 166 genes (13%, Table S1) were also listed in another ORF57-CLIP study using a different anti-ORF57 antibody and a different experimental approach (44). Among of them, 821 ORF57 BS (Table S2) were predominantly mapped to the coding (CDS) (68.6%) and intron (18.3%) regions of the RNA transcripts of 544 unique protein-coding genes (Fig. 1C and 1D). To gain insights into the molecular functions of ORF57-bound RNAs, we performed a gene ontology (GO) analysis by g:Profiler (56) (https://biit.cs.ut.ee/gprofiler/gost) using terms for molecular function (MF), biological processes (BP) and cellular components (CC). We found that ORF57 binds preferentially to the RNAs of which their encoded proteins are involved in the mRNA metabolism and RNA biology (Fig. 1E-1F). Although our anti-ORF57 CLIP-seq also identified ORF57-RNA interactions in the KSHV transcriptome, this report will only focus to novel ORF57-bound host protein-coding mRNAs.

### ORF57 regulates the expression of a subset of host genes during KSHV lytic infection

In parallel by RNA-seq, we compared the host gene expression profiles of BCBL- 1 cells with KSHV latent infection lacking ORF57 expression over the KSHV lytic infection with abundant ORF57 expression (Fig. S1A), along with HEK293T cells ectopically expressing ORF57 over the cells with no ORF57 expression (Fig. S1B). We identified 1,264 up-regulated and 851 down-regulated genes in BCBL-1 cells with KSHV lytic replication and 299 up-regulated and 37 down-regulated genes in HEK293T expressing ORF57 by two-fold changes (p ≤ 0.001, FDR ≤ 0.05) (Fig. S1C-D, Table S3 and S4).

To identify host coding genes in BCBL-1 cells directly regulated by ORF57, we overlapped the differentially expressed host genes as determined by the RNA-seq data described above to 544 genes containing the ORF57 BS identified by ORF57 CLIP-seq. By further selection of highly expressed (RPKM ≥3) protein-coding genes, we found that 20 genes with the ORF57 BS were differentially expressed in BCBL-1 cells with lytic KSHV infection. Of those, 13 were upregulated and 7 downregulated (Fig. 1G and 1H). RNA reads-coverage maps of FOS, PMF1, and LY9 with the ORF57 BS and RBL without an ORF57 BS were selectively illustrated by IGV (Fig. 2A). In HEK293T cells expressing ORF57, we identified four genes with the ORF57 BS as being upregulated, but only two of them were consistently upregulated in both BCBL-1 and HEK293T cells upon ORF57 expression (Fig 1G and 1H). They were FOS, an important component of heterodimeric transcription factor AP-1 for transactivating expression of many genes (5, 6), and CITED2, an inhibitor blocking transactivation of HIF1A-induced genes (57, 58). We also identified additional 7 genes (CDKN2B, DHRS2, RGS2, SERPINI1, FAS, SAT1, and RBL1) lacking the ORF57 BS being differentially expressed, but probably indirect to ORF57, both in BCBL-1 cells with KSHV lytic infection and in HEK293T cells expressing ORF57 (Fig. 1G and 1H).

**Figure 2.**
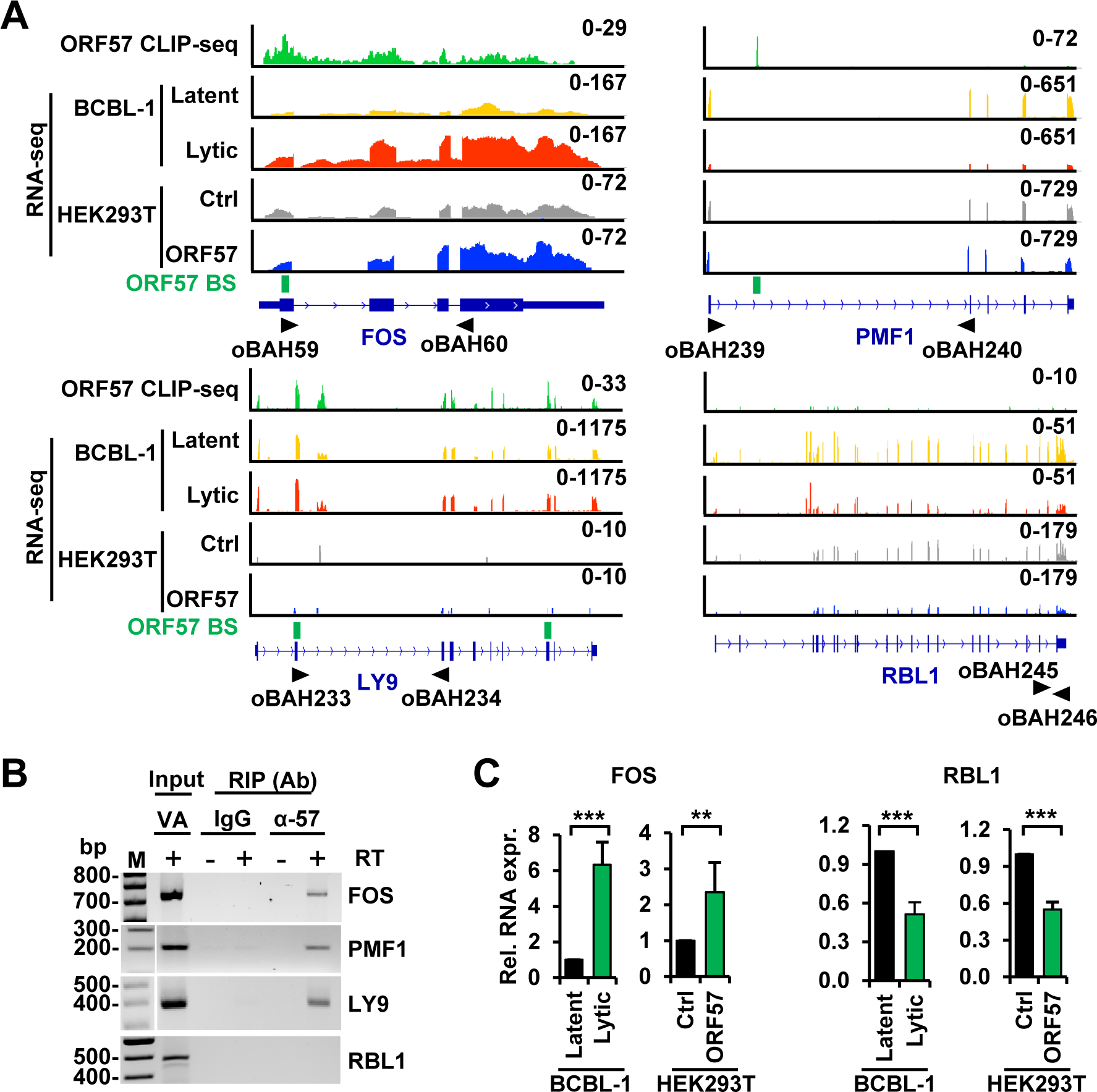
ORF57 binds to target RNAs to regulate the expression of host protein-coding genes. (A) Distribution of ORF57 CLIP-seq reads (on the top in green) obtained from BCBL-1 cells with lytic infection along with the RNA-seq reads-distribution for the longest coding RNA isoforms of FOS, PMF1, LY9, and RBL1 genes in BCBL-1 cells with latent (yellow) or lytic (red) infection and in HEK293T cells transfected with a control empty (Ctrl, gray) or ORF57-expressing (ORF57, blue) vector using the Integrative Genomics Viewer (IGV). The y-axis represents reads-count scale with reads-coverage depth for each sample shown in the upper right corner. The named black arrows below mark the locations of the primers used in RT-PCR detections in (B). The ORF57-binding sites (BS, green box) identified by Piranha software are also shown. RBL1 lacking ORF57 BS by Piranha software is included for the comparison. (B) Validation of ORF57 interacting target RNAs from (A) by RIP-RT-PCR. RT-PCR was performed in the presence (+) or absence (-) of RT using the RNAs isolated from ORF57 RIPs. IgG served as a negative RIP control. Total RNA from BCBL-1 cells induced with VA (1 mM) for 24 h was used as an input control. (C) The effect of ORF57 on FOS and RBL1 RNA expression was determined by RT- qPCR using total RNA from BCBL-1 cells with latent or lytic infection or HEK293T cells transfected with an empty (control Ctrl) or ORF57-expressing vector. GAPDH served as an internal control. Data presented as mean ± SD from three replicates in one representative of three separate experiments. **p < 0.01; ***p < 0.001 by two-tailed Student *t*-test.

By RNA immunoprecipitation (RIP), we subsequently confirmed ORF57 binding to FOS, PMF1, LY9, SOCS3, EMP3, and FASN RNAs, but not to RBL1 in BCBL-1 cells with lytic KSHV infection (Fig. 2B, Fig. S1E). RT-qPCR further confirmed that ORF57 mediated upregulation of FOS, but downregulation of RBL1 both in BCBL-1 cells with lytic KSHV infection and in HEK293T cells with ORF57 expression (Fig. 2C). Together, our data indicate that ORF57 selectively regulates the expression of host genes mainly via direct interaction with the regulated RNAs but only a small fraction of the affected genes indirectly through other host factors.

Considering that FOS is a well-known proto-oncoprotein and a heterodimer component of transcription factor AP-1 (5, 6) previously identified to be upregulated during KSHV lytic infection (12, 15), we next focused on how ORF57 regulates FOS expression and the functional role of elevated FOS expression during KSHV lytic infection.

### ORF57 is essential for FOS expression during KSHV lytic infection

Given the significant increase of FOS RNA in KSHV lytic infection and HEK293T cells with ORF57 expression (Fig. 2A and 2C), we further demonstrated the increased expression of FOS protein by Western blot in these two types of cells under the same conditions (Fig. 3A). To determine whether ORF57 alone is responsible for FOS upregulation during KSHV lytic infection, we evaluated FOS protein expression in iSLK/Bac16 cells harboring a wild-type (wt) or an ORF57 knockout (57KO) KSHV genome (17). As shown in Fig. 3B and when compared to the cells carrying a wt KSHV genome, loss of ORF57 expression from the 57KO genome in iSLK/Bac16 cells dramatically reduced FOS protein expression during KSHV lytic replication, despite only a slight reduction of RTA expression as an indicator of KSHV reactivation (Fig. 3B). To confirm the dependence of FOS expression on ORF57, we transduced the 57KO iSLK/Bac16 cells with a lentivirus expressing wt ORF57. The rescued ORF57 expression in the 57KO cells was found to significantly restore the expression of both FOS RNA (Fig. 3C) and protein (Fig. 3D) as well as RTA protein level (Fig. 3D) after 48 h of KSHV lytic induction. Together, these results indicate that the high expression level of FOS is strongly and uniquely dependent on viral ORF57 during KSHV lytic infection.

**Figure 3.**
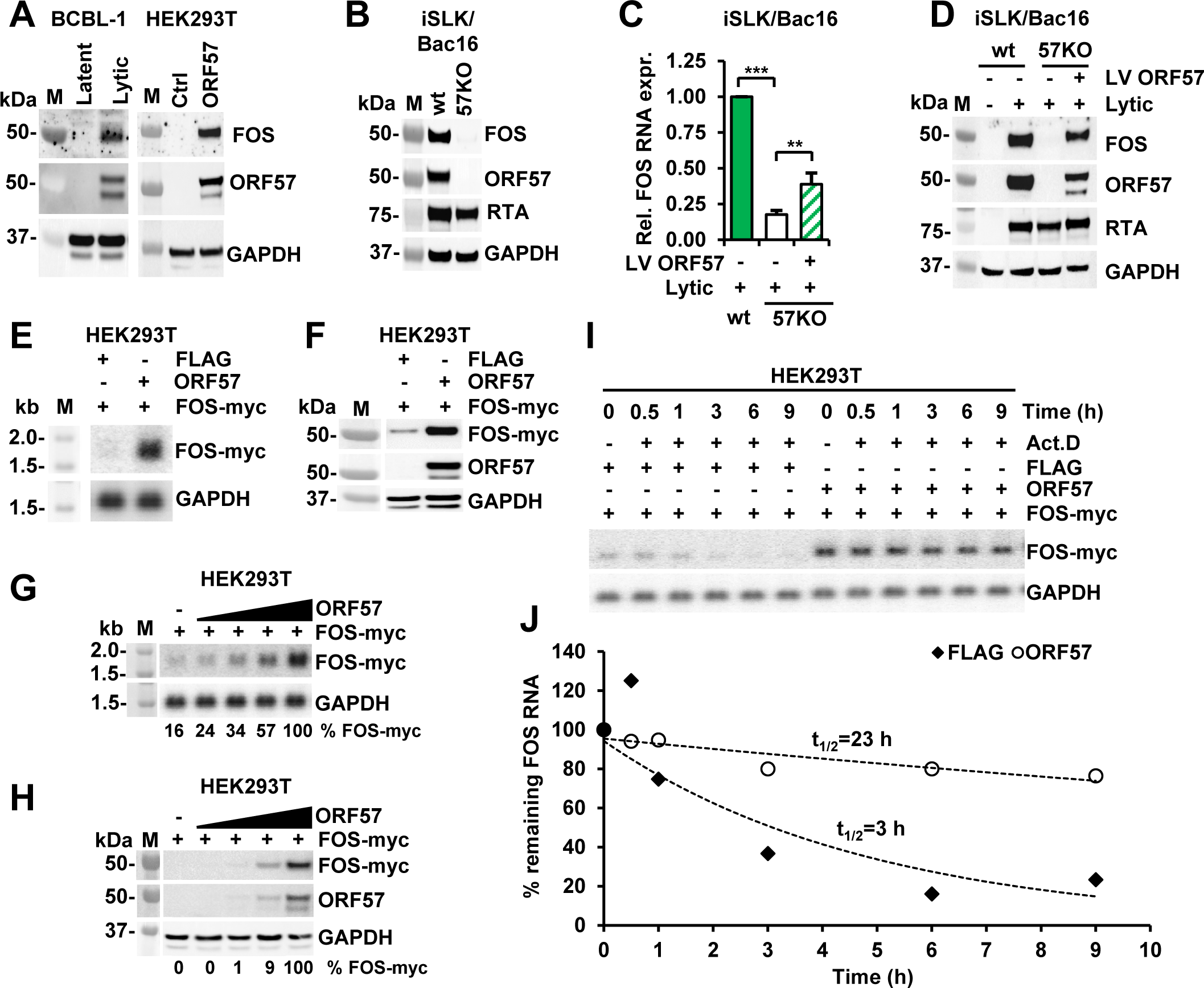
ORF57 is essential for upregulation of FOS expression by stabilizing FOS RNA. (A) ORF57 promotes expression of endogenous FOS protein in BCBL-1 and HEK293T cells. Total cell lysate from BCBL-1 cells with KSHV latent or lytic infection or from HEK293T cells transfected with an empty vector (ctrl) or ORF57-expressing vector (ORF57) was blotted with the corresponding antibodies. Host GAPDH served as a protein loading control. (B) Loss of FOS expression in lytic KSHV infection in iSLK/Bac16 cells containing an ORF57-knockout (57KO) KSHV genome generated by CRISPR-Cas9 technology. Western blot analysis of total cell lysate from the wild-type (wt) and 57KO iSLK/Bac16 cells with KSHV lytic induction for 48 h was blotted for expression of FOS, ORF57, RTA (a marker of KSHV activation), and GAPDH (loading control) proteins. (C and D) Restoration of ORF57 expression in the 57KO iSLK/Bac16 cells rescues the expression of endogenous FOS RNA (C) and protein (D). The 57KO iSLK/Bac16 cells were transduced by ORF57-expressing lentiviruses (LV-ORF57) at MOI 10 followed by induction of KSHV lytic replication by the treatment with sodium butyrate (1 mM) and doxycycline (1 μg/ml) for 48 h before collection of total cell RNA for RT-qPCR (C) and total cell lysate for Western blotting (D). M, protein markers. Data (C) presented as mean ± SD from three replicates in one representative of three separate experiments. **p < 0.01, ***p < 0.001 in two-tailed Student *t-*test. (E and F) ORF57 promotes the expression of exogenous FOS-myc RNA and protein in HEK293T cells. The cells were co-transfected with a FOS-myc expression vector (pCMV6-FOS, 100 ng) together with an ORF57-expression vector (pVM7, 300 ng) or empty plasmid (FLAG, 300 ng). FOS-myc RNA and protein levels at 24 h of co-transfection were determined by Northern blot (E) and Western blot (F), respectively. GAPDH RNA (E) and protein (F) served as a loading control in each assay. (G and H) ORF57 promotes exogenous FOS expression in a dose-dependent manner. HEK293T cells were co-transfected with 100 ng of a FOS-myc plasmid together with 0, 20, 50, 100, or 150 ng of an ORF57-expressing (ORF57) vector. Northern blot (G) and Western blot (H) analyses at 24 h were performed to detect FOS-myc RNA and protein. GAPDH RNA (G) and protein (H) served as a loading control in each assay. Densitometry analysis of individual RNA and protein bands was performed and % FOS- myc RNA (G) or protein (H) in each sample was calculated after normalization to GAPDH. (I and J) ORF57 stabilizes FOS RNA by prolonging FOS RNA half-life in a transcription pulse-chase assay. HEK293T cells were co-transfected with a FOS-myc expressing vector along with an empty FLAG control vector or an ORF57 expressing vector. Twenty-four hours after transfection the RNA polymerase II inhibitor, actinomycin D (Act. D) was added to stop transcription and total RNA was collected at the indicated time up to 9 h. The amount of remaining FOS-myc RNA was determined by Northern blot (I). FOS RNA level in each sample measured by densitometry was normalized to GAPDH and plotted (J) to generated RNA decay curve using the FOS-myc RNA level in each group at 0 h time (t = 0, Act. D adding time) as 100%. Data were one representative of three independent experiments.

### ORF57 stabilizes FOS RNA to promote FOS expression in a dose-dependent manner

FOS expression is regulated at the transcriptional, posttranscriptional, and translational levels (11). ORF57 promotes the expression of its targets primarily at the posttranscriptional level by promoting RNA stability, splicing, and translation (16). To determine the mechanism of FOS regulation by ORF57, we systematically evaluated all levels of FOS expression regulation. First, we found no changes in FOS RNA splicing in BCBL-1 cells from KSHV latent to lytic infection and in HEK293T cells with ORF57 expression versus the without by comparative analyses of the RNA-seq reads-distribution, splice junction reads, and RT-PCR. Then, we transiently transfected HEK293T cells with a myc-tagged FOS cDNA expression vector under control of a HCMV IE promoter with or without co-transfection of an ORF57 expression vector. We found that both FOS RNA (Fig. 3E) and protein (Fig. 3F) were upregulated by ORF57 expression, indicating that ORF57 regulates FOS expression independently of FOS promoter and RNA splicing. Subsequently, we evaluated FOS RNA and protein expression in the presence of increasing amounts of ORF57. We observed a positive correlation of FOS RNA (Fig. 3G) and protein (Fig. 3H) to ORF57 protein levels. Together, these data indicate that ORF57 regulates FOS expression after FOS RNA transcription and splicing in a dose-dependent manner.

ORF57 was previously shown to promote the expression of several viral targets by enhancing the stability of their RNAs, resulting in the accumulation of viral target RNAs in the presence of ORF57 (28, 32, 34, 36, 37, 39). To test if ORF57 also stabilizes FOS RNA, we performed transcriptional pulse-chase experiments in HEK293T cells co-transfected with myc-FOS along with an ORF57-expressing or an empty FLAG control vector. Transcription by RNA polymerase II was inhibited by actinomycin D treatment and RNA samples were collected at various time points. The level of FOS RNA at each time point was determined by Northern blot and normalized to GAPDH RNA levels (Fig. 3I). The values expressed as the percentage of remaining FOS RNA, with levels at 0 h being 100%, were plotted. An exponential regression curve was fitted to the obtained data points to determine FOS RNA half-life (Fig. 3J). We found that in the absence of ORF57, FOS RNA half-life time in the FLAG vector control was ∼3 h; however, its RNA half-life was considerably increased to ∼23 h in the presence of ORF57 (Fig. 3J).

Finally, we performed cell-free *in vitro* translation assays using rabbit reticulocyte lysates in the presence of ^35^S-methionine to determine a possible FOS regulation at the translational level by ORF57. FOS RNA was transcribed under T7 promoter *in vitro* and used as a template for *in vitro* translation in the presence of increasing concentrations of purified recombinant ORF57 protein. We did not observe any effect of ORF57 on FOS protein translation at any tested concentration of ORF57 protein (Fig. S2). Together these data indicate that ORF57 regulates FOS expression at the posttranscriptional level by stabilizing FOS RNA.

### An ORF57-binding motif in FOS RNA is necessary for FOS RNA stabilization

To map the interaction between ORF57 and FOS RNA, we performed a protein pulldown assay using a series of biotinylated short RNA oligomers (oBAH108, oBAH109, oBAH110, oBAH111) covering the entire 59-nt ORF57 BS of FOS RNA identified by Piranha software in the ORF57 CLIP-seq (Fig. 4A). The total protein obtained from HEK293T cells expressing ORF57 was used as a source of ORF57 protein. The previously identified ORF57-binding RNA oligo (oNP42) and −non-binding control RNA oligo (oNP41), derived from vIL-6 RNA, served, respectively, as a positive and negative control (29, 36). We revealed that ORF57 binds efficiently to two partially overlapping oligonucleotides, oBAH109 and oBAH110, derived from the central region of the ORF57 BS identified by ORF57 CLIP-seq (Fig. 4B). Based on these data, we identified the 13-nt sequence 5ʹ-UGCAGCAGCGCGU-3ʹ as the ORF57-binding core motif in FOS RNA. To measure the contribution of the identified binding core motif to overall ORF57 binding to FOS RNA, we replaced the core motif on oBAH110 oligo with 13-nt 5ʹ- GCUUCUGACGAAU-3ʹ sequence from the negative control oligo (oNP41) that lacks ORF57-binding activity. The resulted RNA oligo, oBAH156 with 11-nt mutations (the underlined nts) from the oBAH110 (Fig. 4A), exhibited a 50% reduction of ORF57 binding when compared to wt oBAH110 oligo (Fig. 4C), demonstrating that the 5ʹ- UGCAGCAGCGCGU-3ʹ core motif is mainly responsible for ORF57 binding to FOS RNA.

**Figure 4.**
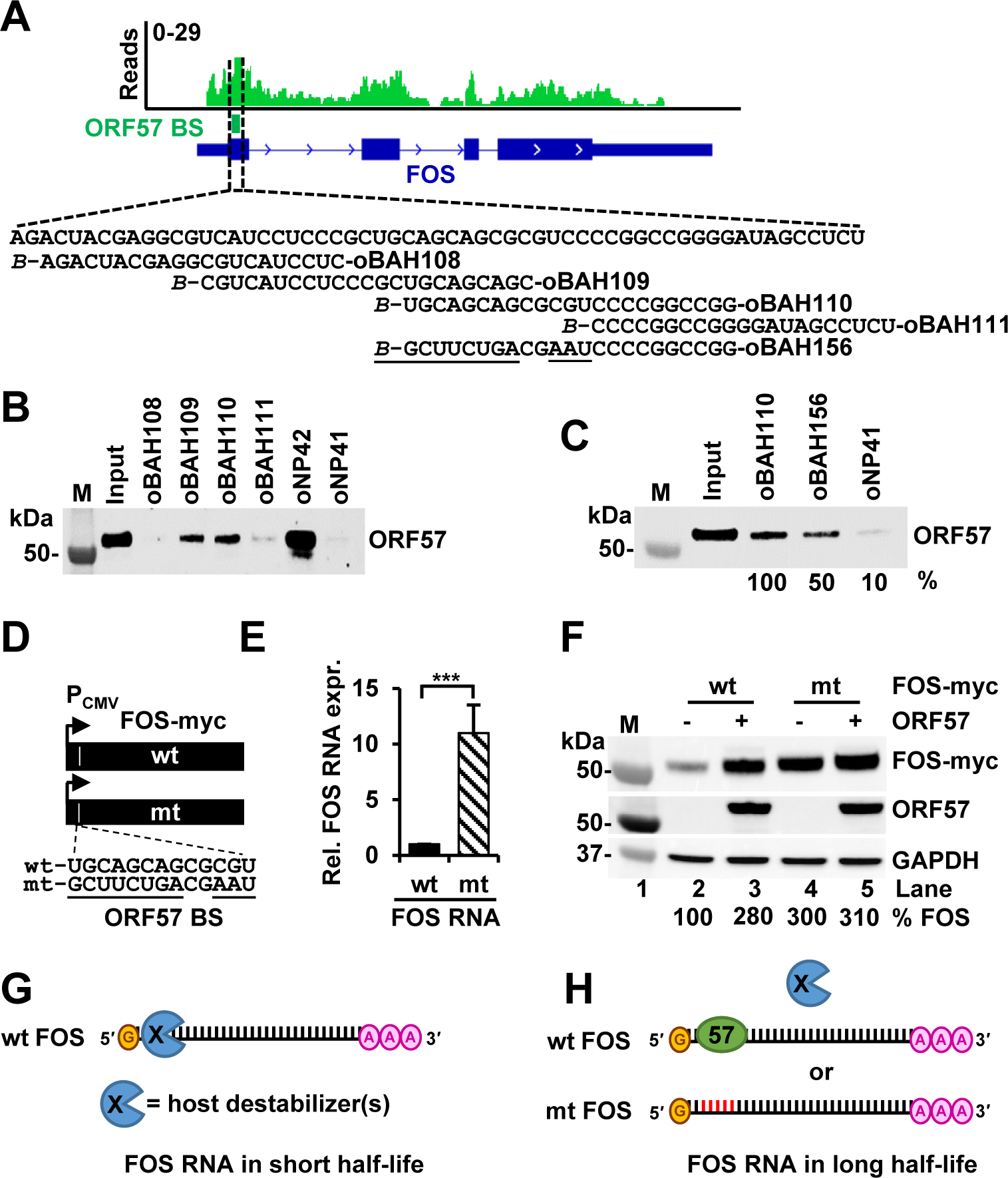
An ORF57-binding motif in FOS RNA is necessary for FOS stabilization. (A) IGV visualization of ORF57 CLIP-seq reads-distribution on FOS RNA, with an ORF57-BS identified by Piranha software and marked by a dashed box. Below is the ORF57 BS nucleotide sequence in FOS RNA and biotinylated (*B-*) RNA oligomers derived from this region used for RNA pulldown assays. (B-C) Identification of an ORF57 BS core in FOS RNA for ORF57 binding by RNA oligo pulldown assays, with KSHV vIL6-derived oNP42 serving as a positive and oNP41 as a negative ORF57-binding control (29). The amount of ORF57 in each oligo pulldown was determined by Western blot using anti-ORF57 antibody. Mutant sequences in oBAH156 are underlined in the panel A. (D) Diagrams of wild-type (wt) and mutant (mt) ORF57 BS in FOS-myc expression vectors and their nucleotide sequences shown below with mutated nucleotides being underlined. (E) Disruption of ORF57 BS by point mutations enhanced FOS RNA expression. FOS RNA levels expressed from the wt and mt FOS-myc vectors (D) after transfection of HEK293T cells for 24 h were quantified by RT-qPCR. GAPDH RNA was used for normalization. Data (E) presented as the mean ± SD from three replicates in one representative of three separate experiments. ***p < 0.001 by two-tailed Student *t*-test. (F) Disruption of ORF57 BS by point mutations enhanced FOS protein expression in the absence of ORF57. FOS-myc protein expression from the wt or mt FOS-myc expression vector (D) was examined after 24 h transfection of HEK293T cells in the presence (+) or absence (-) of ORF57. Total cell lysate was blotted for FOS-myc and ORF57 using the indicated antibodies. GAPDH served as a protein loading control. (G-H) A proposed model of how FOS RNA is stabilized by ORF57 through the ORF57 BS in possible competition with an unknow (X) host destabilizer. In the absence of ORF57, a host RNA destabilizer (s) binds the same ORF57 BS and destabilizes FOS RNA (G), thus making FOS RNA in short half-life. However, in the presence of ORF57 or viral lytic infection, ORF57 binds FOS RNA through the mapped ORF57 BS and prevents the host RNA destabilizer (s) binding to the FOS RNA, thereby, prolonging ORF57 RNA half-life. The same is true when the ORF57 BS is mutated (red) and loss of binding by the host RNA destabilizer (s) or ORF57 increases FOS RNA half-life. FOS RNA 5′ cap (yellow) and 3′ polyA tail (pink A) are also shown in the diagram.

Next, we introduced the same 11-nt mutations into the ORF57-binding core motif of the full-length FOS RNA (Fig. 4D) and evaluated the wt or mt FOS RNA expression in HEK293T cells by RT-qPCR. Surprisingly, we observed an ∼11-fold increase in expression of the mt FOS RNA over the wt FOS RNA (Fig. 4E). This increase led to a 3- fold increase of mt FOS protein expression in the absence of ORF57 (Fig. 4F, compare lanes 4 to 2) and thereby, appeared almost not responding to ORF57 (Fig. 4F, compare lanes 4 to 5).

Based on these results, we hypothesized that, in the absence of ORF57 or during KSHV latent infection, FOS RNA is destabilized by a host destabilizer (s) and has a short half-life (Fig. 4G). Induction of ORF57 expression in BCBL-1 cells during KSHV lytic infection or ectopic ORF57 expression of ORF57 in HEK293T cells leads to ORF57 protein binding to the ORF57-binding core motif within the FOS RNA 5ʹ end to prevent FOS RNA from degradation mediated by the host destabilizer (s) and thus, resulting in increase of FOS RNA half-life (Fig. 4H). Introduction of point mutations into the ORF57 BS not only disrupts ORF57 binding, but also the binding of the host destabilizer (s) to FOS RNA, thereby, promoting FOS RNA accumulation independent of ORF57 (Fig. 4H). Alternatively, ORF57 expression may lead to reduction of expression levels of the putative host destabilizer (s) responsible for FOS RNA instability.

### KSHV reduces cellular RNA nuclease AEN, but not exosomal RNA helicase MTR4, by viral ORF57 via FOS

By analyzing the differentially expressed genes in BCBL-1 cells with lytic KSHV infection (Table S3) and HEK293T cells expressing ORF57 (Table S4), we identified the AEN (apoptosis-enhancing nuclease, also referred as ISG20L1) which bears RNA exonuclease activities (49, 51), as one of the few genes downregulated by ORF57 in both KSHV lytically infected BCBL-1 and ORF57-expressing HEK293T cells (Fig. 5A). The RT- qPCR confirmed AEN downregulation in both cell lines in the presence of ORF57 (Fig. 5B). Further Western blot analysis confirmed ∼45% reduction of AEN protein expression by viral ORF57 in HEK293T cells accompanied by upregulation of endogenous FOS protein level (Fig. 5C).

**Figure 5.**
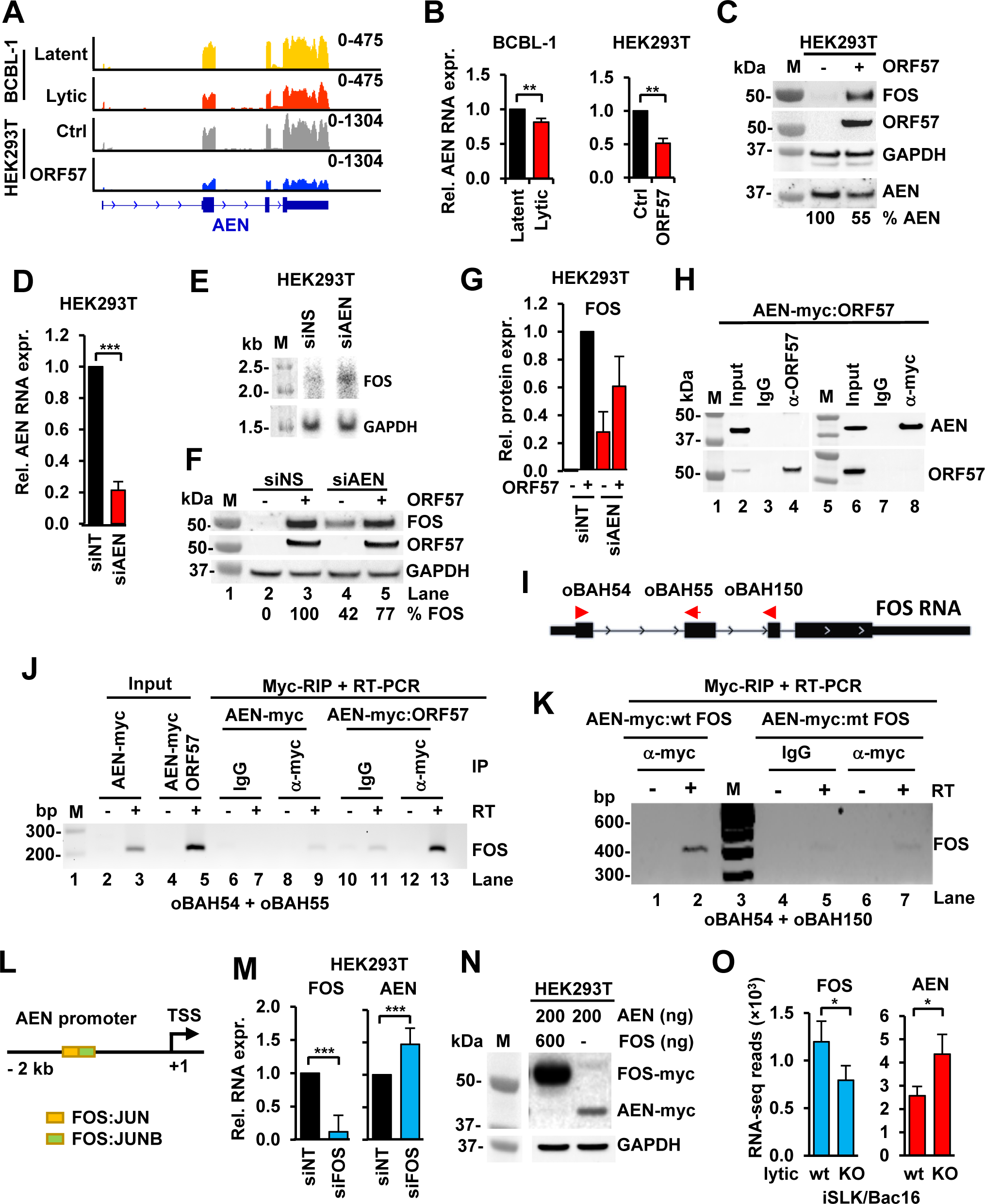
KSHV infection and viral ORF57 upregulate FOS by inhibition of AEN expression to stabilize FOS RNA. (A) RNA-seq reads-coverage showing downregulation of AEN expression in BCBL-1 cells with KSHV lytic infection and in HEK293T cells with ORF57 expression. HEK293T cells transfected with an empty vector served as a control (ctrl). (B) RT-qPCR validation of AEN RNA levels in BCBL-1 cells with KSHV lytic vs latent infection and in HEK293T cells with or without ORF57 expression. Data presented as the mean ± SD from three separate experiments, each with three replicates. ** p < 0.01, in two-tailed Student *t-*test. (C) ORF57-enhanced FOS expression led to reduction of AEN expression in HEK293T cells. Total cell lysate extracted from HEK293T cells without (-) or with (+) expression of ORF57 was blotted for individual endogenous proteins using the corresponding antibodies, with GAPDH serving as a loading control. Protein band densitometry assays in one representative of two repeats were performed and % AEN protein level in each sample was calculated after normalization to GAPDH, with the AEN band intensity in the cells without ORF57 expression set as 100%. (D and E) AEN destabilizes FOS RNA in HEK293T cells. After 48 h of siRNA-mediated AEN knockdown (siAEN), total RNA extracted from HEK293T cells was examined for AEN knockdown efficiency by RT-qPCR (D) and expression of endogenous FOS RNA by Northern blot (E) using a FOS-specific, ^32^P-labeled antisense oligo probe oBAH55 (see panel I). The cells treated with a nontargeting siRNA (siNT) served as a control. GAPDH served as a sample RNA loading control. Data (D) are the mean ± SD from two separate experiments, each with three replicates. ***p < 0.001 by two-tailed Student *t-* test. One representative (E) of two Northern blot repeats is shown. (F and G) Knockdown of AEN expression enhances FOS protein expression in HEK293T cells. After 48 h of siAEN or siNT knockdown, HEK293T cells were then transfected with (+) or without (-) an ORF57 expression vector for additional 24 h and blotted for the indicated protein expression using the corresponding antibodies (F). GAPDH served as a protein loading control. See AEN knockdown efficiency in the panel D. Relative FOS protein expression (G) in siAEN or control siNT-treated cells in the presence or absence of ORF57 were measured as the mean ± SD from two separate experiments. Protein band densitometry were performed and % FOS protein level in each sample was calculated after being normalized to GAPDH, with the FOS band intensity in the cells treated with control siNT setting as 100%. (H) ORF57 protein does not interact with AEN protein in HEK293T cells by co-immunoprecipitation (co-IP) assays. Total cell lysate from HEK293T cells co-transfected with ORF57 and AEN-FLAG-myc expressing vectors for 24 h was co-IPed with an anti-ORF57 or Myc-antibody. ORF57 and AEN-myc proteins from each IP pulldown were blotted with the corresponding antibodies. (I and J) AEN binds endogenous FOS RNA in cells by anti-Myc RNA immunoprecipitations (RIP). (I) Diagram of FOS RNA structure and oligos (red arrows) used for RT-PCR detection of RIPed FOS RNA. (J) HEK293T cells ectopically expressing AEN-myc (pCMV6-AEN) alone or in co-expression with ORF57 were UV- crosslinked for anti-myc RIP of the AEN-RNA complexes. The RNA in the complexes were extracted for RT-PCR detection of FOS RNA in the absence (-) or presence (+) of reverse transcriptase (RT) with a pair of FOS-specific primers oBAH54 and oBAH55 (I). IgG served as a negative RIP antibody control. (K) AEN binds wt, but not mt FOS RNA ectopically expressed in cells by RIP. HEK293T cells co-expressing AEN-myc (pCMV6-AEN) and wt or mt FOS vectors were UV- crosslinked for anti-myc RIP of the AEN-RNA complexes. The RNA in the complexes were extracted for RT-PCR detection of FOS RNA in the absence (-) or presence (+) of reverse transcriptase (RT) with a pair of FOS-specific primers oBAH54 and oBAH150. (L and M) FOS binding to the *AEN* promoter blocks AEN transcription. (L) Diagram shows the *AEN* promoter region (−2 kb) upstream of the AEN transcription start site (TSS) as illustrated in the UCSC Genome Browser. The colored boxes represent the binding sites of FOS:JUN (yellow) and negative regulator FOS:JUNB (green) predicted by JASPAR CORE 2022 - Predicted Transcription Factor Binding Sites (MA1126.1) prediction. (M) Knockdown of FOS expression by siRNA (siFOS) in HEK293T cells, with siNT as a control, increased AEN RNA expression as quantified by RT-qPCR after being normalized to GAPDH RNA. The left panel shows the FOS knockdown efficiency. Data presented as the mean ± SD from all three separate experiments, each with three replicates. ***, p < 0.001 in two-tailed Student *t-*test. (N) FOS inhibits AEN expression in cells. HEK293T cells in six-well plates were co-transfected with the indicated FOS-myc (pCMV6-FOS) and AEN-myc expression vectors (pCMV6-AEN). FOS-myc and AEN-myc protein expression levels at 24 h after the co-transfection were determined by Western blotting with anti-myc antibody. GAPDH served as a sample loading control. (O) Effect of ORF57 KO in KSHV genome on FOS and AEN expression in iSLK/Bac16 cells. Total RNA-seq reads from FOS to AEN displayed an opposite expression profile in iSLK/Bac16 cells containing a wt or 57KO KSHV genome during lytic infection. *, p < 0.05 in two-tailed Student *t-*test.

To determine whether AEN alone does control FOS expression, we knocked down AEN expression (Fig. 5D) in HEK293T cells using an AEN-specific siRNA (siAEN) and demonstrated that knockdown (KD) of AEN expression stabilized FOS RNA, thus increasing endogenous FOS RNA level as determined by Northern blot (Fig. 5E), when compared to the control cells with a non-specific siRNA (siNS) treatment. The cells with AEN KD were also transfected with or without an ORF57-expressing vector and showed ∼42% increase of endogenous FOS protein in the absence of ORF57 when compared to the siNS control cells expressing ORF57 (Fig. 5F, compare lanes 4 to 3, and Fig. 5G), suggesting a fraction of FOS expression in association with AEN. We next examined if the downregulation of AEN protein by ORF57 (Fig. 5C) was resulted from its interaction with ORF57 or indirectly through ORF57-mediated increase of FOS. We first performed a co-immunoprecipitation assay using total protein extracts from HEK293T cells expressing ORF57 and myc-tagged AEN proteins using anti-ORF57 or anti-Myc antibodies, but we did not find any direct protein-protein interaction between AEN and ORF57 (Fig. 5H). By RNA immunoprecipitation (RIP) in combination with RT-PCR of total RNA extracted from the co-transfected cells (Fig. 5I-K), however, we did find that AEN binding to endogenous FOS RNA at basal level and the FOS RNA stabilized in the presence of ORF57 (Fig. 5J, compare lanes 9 to 7 and 13 to 11). Further studies showed that AEN binds to ectopically expressed FOS RNA, but not much so (Fig. 5K, compare lanes 2 to 7) to the ORF57 binding site-mutated FOS RNAs (Fig. 4D). Data indicate that AEN-binding to FOS RNA leads to the instability of FOS RNA and consistently, knocking down AEN expression led to increased expression of FOS protein in the absence of ORF57 (Fig. 5F, compare lanes 4 to 2).

To understand how an increased expression of FOS could result in the decreased expression of AEN (Fig. 5C) and to identify possible regulatory circuit between FOS as a transcription factor and AEN as an RNA nuclease, we analyzed the *AEN* promoter region for transcription factor binding sites by JASPAR CORE prediction program (59) and found a distal enhancer in the *AEN* promoter containing a FOS:JUN binding site followed by a FOS:JUNB binding site immediately downstream (Fig. 5L, diagram). FOS:JUNB binding to a group of promoters was shown to inhibit their activation (60). Accordingly, we found that knockdown of FOS expression (Fig. 5M, left panel) in HEK293T cells led to significant increase of AEN expression (Fig. 5M, right panel). In contrast, ectopic overexpression of FOS was also found to suppress expression of ectopic AEN from a human CMV IE promoter which contains two AP-1 binding sites (61, 62) (Fig. 5N). Consistently, we found that the reduced FOS expression was accompanied by the increased expression of AEN from the 57KO iSLK/Bac16 cells when compared with the wt iSLK/Bac16 cells with lytic KSHV infection (Fig. 5O). Altogether, these data indicate the presence of a mutual regulation or a feedback regulatory mechanism of AEN and FOS expression in the cells that could be interrupted by expression of viral ORF57 either ectopically or by KSHV lytic infection.

It was reported that ORF57 protects viral transcripts likely by preventing the recruitment of MTR4, another cellular RNA helicase and a component of RNA exosome complex linked to nuclear RNA decay (63-66). To test if MTR4 could also function as a host FOS RNA destabilizer, we knocked down MTR4 in HEK293T cells and determined the level of FOS protein by Western blot. As expected, MTR4 knockdown also led to increase of FOS protein level (Fig. S3A). However, MTR4 expression was not significantly altered in BCBL-1 cells during KSHV lytic infection, nor in HEK293T cells expressing ORF57 (Fig. S3B). Like AEN, MTR4 does not interact with ORF57 (Fig. S3C). Altogether, these data indicate that MTR4 regulation of FOS RNA stability in cells is unrelated to ORF57 expression, nor to KSHV lytic infection, which contrasts with AEN regulation of FOS RNA stability interruptible by viral ORF57 and KSHV lytic infection.

### FOS transactivates RGS2 transcription in BCBL-1 cells during KSHV lytic replication and in HEK293T cells expressing viral ORF57

By binding to JUN, FOS forms a well-known heterodimeric transcription factor AP- 1 (5, 6), whose role in KSHV lytic replication has been a subject of several previous studies. AP-1-binding sites have been found in the promoter of several KSHV lytic genes, including immediately early (RTA, K8, K4.1, ORF19), delayed early (ORF37 and ORF61), and late genes (ORF62) (10, 12, 15).

To identify the other host genes potentially regulated by ORF57-mediated increase of FOS expression during KSHV lytic infection, we performed gene set enrichment analysis (GSEA) (67) using transcription factor targets (TFT) database (https://www.gsea-msigdb.org/) to determine the key transcription factors responsible for the expression changes of host genes associated with KSHV lytic infection in BCBL-1 cells. We found AP-1 being a top key transcription regulator in expression of 69 host genes which were upregulated by KSHV lytic infection. Further analysis revealed that the promoter of all 69 genes contains at least one AP-1-binding motif “NTGASTCAG” in the regions spanning 4 kb centered on their transcription starting sites (Fig. 6A) (Table S5). By overlapping of the 69 FOS targets identified in BCBL-1 cells with the genes upregulated in ORF57-expressing HEK293T cells, we identified *RGS2* (regulator of G protein signaling 2) encoding a GTPase activating protein (GAP) (68) as a putative FOS target commonly regulated by ORF57 and lytic KSHV infection (Fig. 6B).

**Figure 6.**
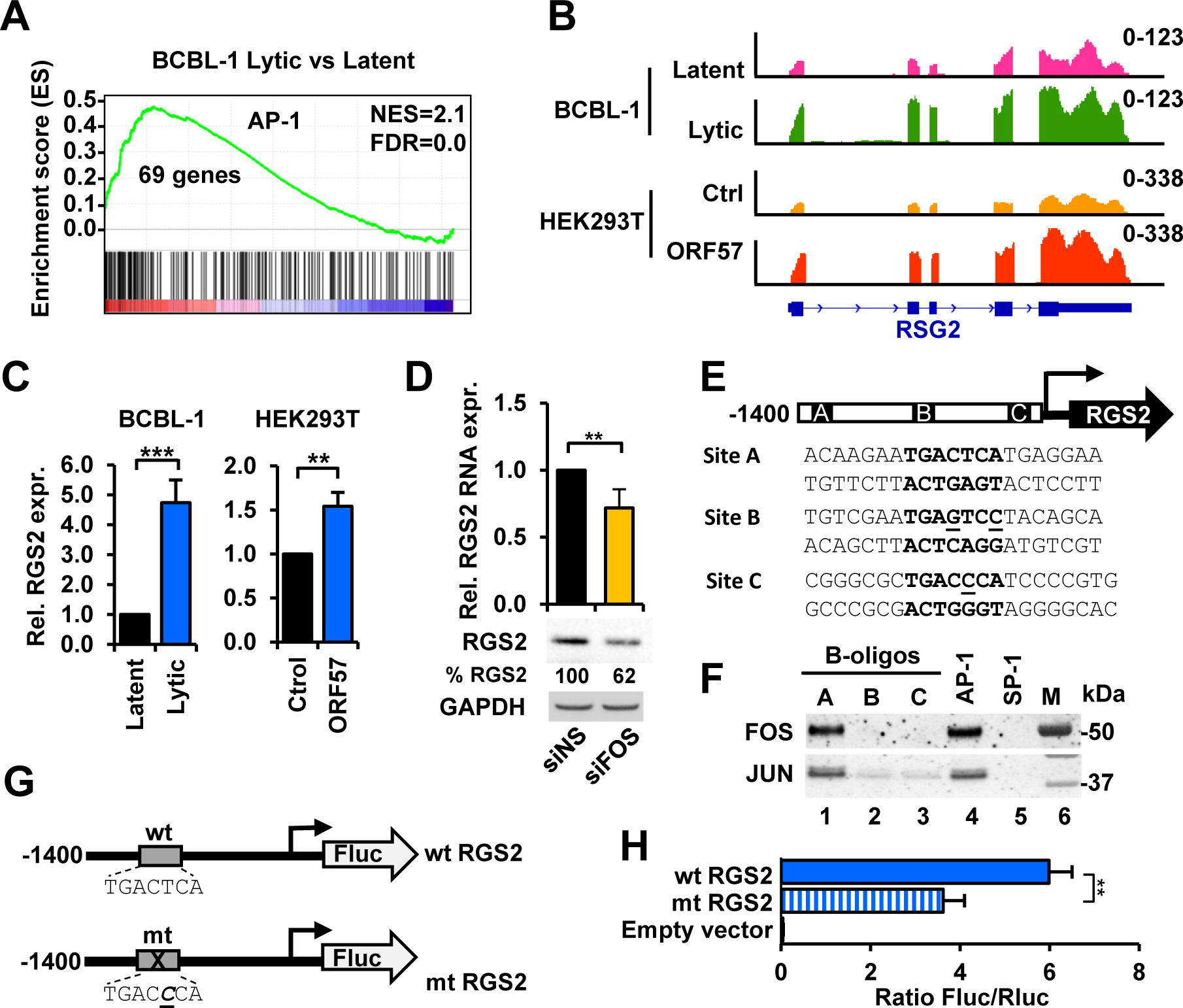
FOS functions as an AP-1 heterodimer component to transactivate *RGS2* expression. (A) Gene set enrichment plot showing AP-1 target genes being selectively enriched in BCBL-1 cells with KSHV lytic infection. (B and C) RNA-seq reads-coverage (B) and verification by RT-qPCR (C) showing the increased expression of RGS2 in BCBL-1 cells 24 h of lytic infection or in HEK293T cells with 24 h of ORF57 expression. Reads-coverage scale at the y-axis (B) represents the number of RNA-seq read counts with a scale in an upper right corner. RT-qPCR data (C) presented as the mean ± SD from three replicates in one representative of three separate experiments. **, p < 0.01; ***, p < 0.001 in two-tailed Student *t-test*. (D) Knockdown of FOS expression by siRNA (siFOS) in HEK293T cells, with a nontargeting siRNA (siNS) as a control, reduced the expression of RGS2 RNA and protein quantified by RT-qPCR (bar graphs) and Western blot (gel blots) after being normalized to GAPDH RNA (RT-qPCR) and protein (gel blots) loading control. RGS2 RNA data presented as the mean ± SD from three separate experiments, each with three replicates. **, p < 0.01 in two-tailed Student *t-*test. (E-H) Identification of an AP-1 BS and its function in the *RGS2* promoter. (E) Predicted AP-1 BS sequences A (−1001 to −981), B (−579 to −559), and C (−147 to −126) upstream of the RGS2 transcription start site (TSS) using JASPAR database. (F) Predicted AP-1 BS A is an authentic AP-1 BS as determined by DNA oligo pull down assays. Biotinylated double-stranded DNA oligos (B-oligos) with the predicted putative AP-1 sequences (E) immobilized on avidin beads were used in oligo pulldown assays with the total cell lysate from HEK293T cells expressing ORF57. DNA oligos containing a consensus AP-1 or SP-1 BS served, respectively, as a positive or negative control. The proteins in the oligo pulldowns were immunoblotted by anti-FOS and anti-JUN antibodies. (G) Diagrams showing the inserted *RGS2* promoter (−1400 to +100 nt relative to TSS) with a wild-type (wt) or mutated (mt) AP-1 BS A upstream of a Firefly luciferase (FLuc) reporter. (H) Introduction of a single nucleotide (T to C) mutation in the RGS2 AP-1 BS A significantly reduced RGS2 protomer activity. Dual luciferase reporter assays using the total cell lysate from HEK293T cells co-transfected with a FOS expression vector (pCMV6-FOS) together with a wt (pBAH8) or mt (pBAH10) RGS2 luciferase reporter for 24 h. A co-transfected Renilla luciferase (Rluc) reporter (pRL-TS) (84) served as an internal control. Data presented as the mean ± SD from three replicates in one representative of three separate experiments. **, p < 0.01 in two-tailed Student *t-*test.

RT-qPCR confirmed the upregulation of RGS2 expression in BCBL-1 cells with lytic replication and in HEK293T cells with ORF57 expression (Fig. 6C). Furthermore, we confirmed the FOS regulation of endogenous RGS2 expression and observed the reduced expression of both RGS2 RNA and protein in HEK293T cells upon knockdown of FOS by a FOS-specific siRNA, siFOS (Fig. 6D). These results suggests that FOS might transactivate *RGS2* transcription by direct binding to its promoter and thus, a partial dependency indirectly on ORF57 expression.

To demonstrate if ORF57 regulates RGS2 expression by FOS transcription activity, we scanned a 1.5-kb region of the *RGS2* locus consisting of 1400 bp upstream and 100 bp downstream of the RGS2 transcription start site (TSS) for the putative FOS or AP-1-binding motifs using the JASPAR database (http://jaspar.genereg.net/). Based on proximity to the TSS and a confident score >7, we selected three (A, B, and C) putative AP-1-binding motifs in the *RGS2* promoter region (Fig. 6E). Biotin-labeled double-stranded DNA oligos corresponding to the putative FOS-binding motifs were used in DNA-protein pulldown assays with HEK293T cell extracts (Fig. 6F). The oligos containing AP-1- and SP-1-consensus binding motifs were used as positive and negative controls, respectively. As shown in Fig. 6F, we found that only oligo A containing a consensus AP- 1-binding motif TGACTCA showed FOS and JUN binding comparable to the AP-1 control oligo. The remaining oligos B and C which miss one (C) or two (B) nucleotides in the AP- 1-binding motif and control SP-1 oligo did not bind either FOS or JUN.

The identified AP-1-binding site in RSG2 regulation was examined by inserting the 1.5-kb DNA fragment from the *RSG2* promoter region into a luciferase reporter. We also constructed a mutant version containing a single nucleotide mutation at −989 T>C in the AP-1-binding motif A (Fig. 6G) as seen in the oligo C binding assay (Fig. 6E-F). The wt and mt promoter activity was then compared by dual-luciferase assay in HEK293T cells as described (36). When compared to the empty control vector, the wt *RSG2* promoter displayed ∼6 fold higher of luciferase activity, while the mt *RGS2* promoter lacking AP-1 binding displayed only 3.6-fold higher of luciferase activity (Fig. 6H), indicating a significant reduction of the *RGS2* promoter activity when a T-to-C point-mutation in the AP-1 A-binding site was introduced.

### FOS and RGS2 are partially responsible for AKT phosphorylation

To further confirm the FOS expression in dependence of ORF57 during KSHV lytic infection (Fig. 3B-D) and transactivation of RGS2 expression being the one of FOS functions (Fig. 6), the iSLK/Bac16 cells with a wt or 57KO KSHV genome were used in the studies shown in Fig. 7A-B. We demonstrated that ORF57 KO in iSLK/Bac16 cells led to reduced RGS2 RNA level during KSHV lytic infection (Fig. 7A). Most importantly, loss of RGS2 RNA expression in the 57KO iSLK/Bac16 cells could be partially rescued by transduction of the cells with an ORF57-expressing lentivirus (Fig. 7A).

**Figure 7.**
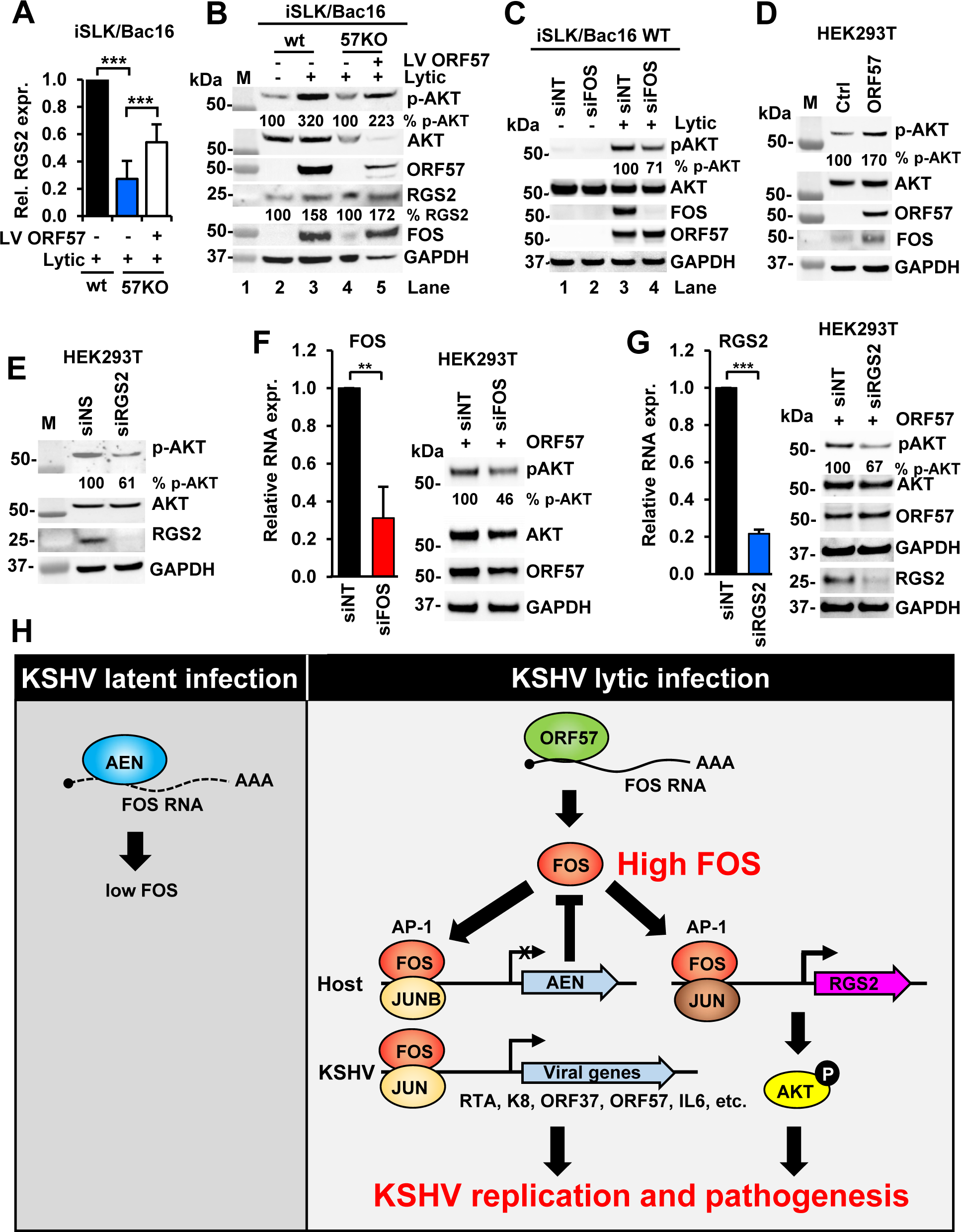
KSHV ORF57-mediated expression of FOS promotes AKT phosphorylation partially through upregulation of RGS2. (A) Reduced RGS2 expression in the iSLK/Bac16 cells containing an ORF57-null KSHV genome (57KO) during KSHV lytic infection could be partially rescued by ORF57- expressing lentivirus (LV ORF57) infection. The iSLK/Bac16 cells with wild-type (wt) KSHV genome during lytic infection served as a control. Total cell RNA extracted from iSLK/Bac16 cells were quantified for RGS2 expression by RT-qPCR. Data presented as the mean ± SD were from three replicates in one representative of three separate experiments. *** p < 0.001 by two-tailed Student *t-*test. (B) Increased RGS2 by FOS triggers AKT phosphorylation (p-AKT) during KSHV lytic infection in iSLK/Bac16 cells. The wt and 57KO iSLK/Bac16 cells without or with transduction of ORF57-expressing lentivirus (LV-ORF57) were induced for KSHV lytic infection for 48 h and followed by Western blot for expression of p-AKT, total AKT, ORF57, RGS2, and FOS, with GAPDH serving as a loading control. (C) Knockdown of FOS expression during KSHV lytic infection in iSLK/Bac16 cells led to decreased phosphorylation of AKT. The wt iSLK/Bac16 cells were treated with a non-targeting (siNT) or FOS-specific (siFOS) siRNA 24 h before induction (+) of lytic replication. The protein samples harvested 48 h after the induction were analyzed in Western blot for expression of p-AKT, total AKT, FOS, ORF57. and GAPDH. Cellular GAPDH served as a sample loading control. The cells without induction (-) served a negative control. Level of p-AKT phosphorylation was normalized to total AKT and GAPDH with the level of p-AKT in siNT-treated cells in KSHV lytic infection being set as 100%. (D and E) Both RGS2 and FOS in HEK293T cells contribute partially to AKT phosphorylation. (D) ORF57-mediated FOS expression promotes AKT phosphorylation. Total cell lysates from HEK293T cells transfected with an empty control (ctrl) or ORF57- expressing (ORF57) vector for 24 h were immunoblotted for p-AKT, total AKT, ORF57, FOS, and GAPDH. (E) Knockdown of RGS2 expression in HEK293T cells led to reduced AKT phosphorylation. Total cell lysates from HEK293T cells treated by siNT or siRGS2 (siRNA-specific for RGS2) at 24 h were immunoblotted for p-AKT, total AKT, RGS2, and GAPDH with the corresponding antibodies. GAPDH protein served as a sample loading control. (F and G) Knockdown of FOS or RGS2 expression in HEK293T cells in the presence of ORF57 led to reduced AKT phosphorylation. HEK293T cells 24 h after ORF57 transfection were treated with siNT or siFOS (F) or siRGS2 (G) for another 24 h and then the collected total cell lysates were used for immunoblotting of p-AKT, total AKT, and ORF57, and GAPDH with the corresponding antibodies. GAPDH protein served as a sample loading control. Knockdown efficiency of FOS (F) and RGS2 (G) RNA quantified by RT-qPCR performed in triplicates are shown in bar graph in each panel. ** p < 0.01; *** p < 0.001 by two-tailed Student *t-*test. RGS2 protein expression in the siRGS2 knockdown cells are also shown in the panel G. Level of p-AKT phosphorylation was normalized to total AKT and GAPDH with the siNT-treated cells expressing ORF57 being set as 100%. (H) A proposed model of mutual regulation or feedback regulatory mechanism of AEN and FOS expression in the cells during KSHV lytic infection. The increased FOS RNA stability and protein expression mediated by ORF57 inhibits AEN expression, but transactivates viral and host gene expression, including host *RGS2* which promotes AKT phosphorylation, and consequently, leading to viral pathogenesis.

Given that a high level of RGS2 activates PI3K/AKT pathway (52), we subsequently evaluated the correlation between RGS2 expression and AKT activation in wt iSLK/Bac16 cells during KSHV lytic infection. We observed a significant increase of AKT phosphorylation (p-AKT) upon the expression of ORF57 along with the increased FOS and RGS2, while total AKT protein remained no change (Fig. 7B, compare lanes 3 to 2). This phenomenon was abolished by ORF57 KO (Fig. 7B, lane 4) but could be rescued by restoring ORF57 expression in the 57KO iSLK/Bac16 cells by an ORF57 lentivirus infection (Fig. 7B, compare lanes 5 to 4). Knockdown of FOS expression in iSLK/Bac16 cells during lytic KSHV infection led to partial reduction of AKT phosphorylation (Fig. 7C, compare lanes 4 to 3).

In HEK293T cells with ORF57 expression, we confirmed that ORF57-induced expression of FOS led to increased level of p-AKT, but not total AKT protein (Fig. 7D). Accordingly, knockdown of RGS2 expression in HEK293T cells also led to reduced level of p-AKT without affecting total AKT expression (Fig. 7E). The knockdown of FOS (Fig. 7F) or RGS2 (Fig. 7G) expression in HEK293T cells in the presence of ORF57 expression further confirmed their roles in partial regulation of AKT phosphorylation. Together, these data conclude that ORF57 promotes AKT phosphorylation by upregulating the expression of FOS and RGS2.

## DISCUSSION

High-throughput RNA-seq and CLIP-seq have been widely used to detect and profile the entire transcriptome and specific protein-RNA interactions in cells. We and others have applied RNA-seq and CLIP-seq technologies to identify viral and host RNAs as ORF57 targets (44, 46, 54, 69). To date, many CLIP-seq technologies have been developed and widely used in screening genome-wide RBP-RNA interactions in living cells (70-72). However, lack of rigorous validations after RNA CLIP-seq in most reports is common. Hypothesis-driven reports purely based on bioinformatics analysis without laboratory experimental verification have hindered our understanding of biological phenomena and have given rise to misleading biological concepts. In this report by applying ORF57 CLIP- seq using an affinity-purified, extremely high specific anti-ORF57 antibody in combination with RNA-seq analyses of BCBL-1 cells with lytic KSHV infection and HEK293T cells with ectopic KSHV ORF57 expression, we identified a subset of host protein-coding RNA transcripts regulated by KSHV posttranscriptional regulator ORF57. This subset partially overlapped with a previous study (44) of which a different anti-ORF57 antibody and a different CLIP-seq approach were used. By rigorous validations with different means which were lacking in the other study (44), we found that FOS and CITED2 RNAs interact with ORF57 and were the two major host targets significantly upregulated in the presence of ORF57.

FOS dimerizes with JUN as a well-known transcription factor AP-1 to transactivate the expression of both host and viral genes (5-7). AP-1 binds to viral promoters of RTA, K8, ORF57, K4.1, ORF19, ORF37, ORF61, ORF62, and IL6 and increases the transcription of both viral and host genes (10, 12, 13, 15). AP-1 or FOS/JUN upregulation during KSHV lytic infection has been reported (10, 12, 13, 15). The mechanism of how FOS/JUN could be upregulated during KSHV lytic infection was previously linked to multiple dysregulated possible cellular activities. These include that KSHV RTA transactivates AP-1 expression and increases AP-1 binding affinity to the responsible promoters (10), KSHV infection increases JUN phosphorylation (10) (12), and viral ORF45 mediates FOS protein phosphorylation and stability through ERK-RSK activation (15), because FOS is a sensor of MAPK activation (12, 13, 15, 73). In this report, we discovered a novel mechanism of FOS upregulation by KSHV ORF57-mediated stability of FOS RNA.

We found that KSHV ORF57 binding FOS RNA via a 13-nt RNA motif near the FOS RNA 5ʹ end prolongs FOS RNA half-life. This binding of ORF57 to FOS RNA appears to competitively prevent a host RNA destabilizer(s) from association to FOS RNA, because mutations introduced into the motif also promotes FOS RNA stability in the absence of ORF57. To seek for such a host factor destabilizing FOS RNA, we discovered AEN, a host apoptosis enhancing nuclease, but not an exosomal RNA helicase MTR4 (63-66), as a sensitive responder of KSHV ORF57 and FOS expression. We found that reduction of AEN expression was inversely correlated to increase of FOS expression in BCBL-1 cells with lytic KSHV infection and HEK293T cells with ectopic ORF57 expression. FOS appears to block AEN transcription. By binding to JUN (also JUNB or JUND) or FOS, FOS (also FOSB, FRA1 or FRA2) forms a well-known heterodimeric or homodimeric transcription factor AP-1 which could be oncogenic or tumor-suppressive depending on the cell type (5, 6). Mechanistically, the cells achieve a high level of FOS expression by directly binding of ORF57 and stabilizing FOS RNA and by FOS-reduced expression of AEN which degrades FOS RNA. The latter is exercised presumably by FOS binding to the *AEN* promoter region and suppressing the *AEN* promoter activity. This presumption is supported by the observation that knockdown of FOS expression led to increase AEN expression. The *AEN* promoter region has two predicted FOS:JUN and FOS:JUNB binding motifs separated by 295 bp of nucleotide sequences. Although FOS binding and point mutation experiments in the *AEN* promoter region are needed in the future studies, the FOS:JUNB had been shown to function as a negative regulator for transcription (60).

RGS (regulator of G protein signaling) family has over 20 members involving in G protein signaling through G protein-coupled receptors (74, 75). RGS2 (76) functions as a GTPase activating protein (GAP) to increase the natural GTPase activity and accelerate the intrinsic GTP hydrolysis activity of Gα subunits (68). RGS2 is a multifunctional regulator important for cardiac and smooth muscle function (77), neuronal plasticity (78), and preeclampsia (79). Recent studies in different cell systems showed that RGS2 attenuates G protein signaling in chemo-resistant non-small cell lung cancer cell lines (80), but promotes AKT phosphorylation in human prostate adenocarcinoma cell line LNCaP cells (52). RGS2 also inhibits or promotes protein translation in a stress-related manner (81-83). In our study, we revealed the increased expression of RGS2 in BCBL-1 cells with KSHV lytic infection and HEK293T cells with ectopic ORF57 expression along with increased level of transcription factor FOS protein. We uncovered that FOS transactivates RGS2 expression by binding to a consensus AP-1 sequence motif in the *RGS2* promoter, resulting in the enhancement of AKT phosphorylation.

In summary and illustrated in Fig. 7H, we discovered that KSHV lytic infection promotes FOS expression through viral ORF57-mediated stabilization of FOS RNA. The resulted high level of cellular FOS in KSHV lytic infection inhibits host nuclease AEN transcription, but transactivates RGS2 expression to activate host AKT/ERK pathway. Together with other studies, we conclude that FOS RNA is stabilized by binding ORF57 during KSHV lytic infection. As AP-1 is a heterodimer of FOS/JUN and binds to KSHV viral promoters of lytic genes and host IL6 to increase the transcription of both viral and host genes (10, 12, 13, 15), a constant high level of oncoprotein FOS expression during KHSV lytic infection would lead to cascaded viral and host gene expression, host cell responses, and an onco-pathogenic outcome.

### Data and code availability

Original CLIP-seq and RNA-seq data have been deposited in NCBI’s Gene Expression Omnibus (NCBI GEO) with the accession numbers (Acc. No.) GSE179726, GSE179727, and GSE179728.

## Acknowledgments

This research was supported by the Intramural Research Program of the National Cancer Institute, Center for Cancer Research, NIH (ZIASC010357 to Z.M.Z.). We thank Xuefeng Liu of Georgetown University for providing us his anti-myc antibody as a gift in this study and Abdul Waheed and Eric Freed of National Cancer Institute for their assistance in ORF57-lentivirus production.

## Author contributions

V.M., B.A., and Z.M.Z. designed the study. V.M., B.A., and Y.M. performed the experiments. V.M., B.A., Y.M., A.L., M.C., and Z.M.Z. analyzed data and participated in the discussion and interpretation of the results. V.M., B.A., and Z.M.Z drafted the manuscript. V.M., B.A., and Z.M.Z. revised the manuscript. All authors read and approved the final manuscript.

## Materials and Methods

### Cell culture

Human renal carcinoma cell line-derived iSLK cells carrying the bacterial artificial chromosome (Bac16) containing the wild type (wt) or CRISPR/Cas9-generated ORF57 knockout (57KO) KSHV genomes (17, 85) and HEK293T, human embryonic kidney cells, were cultured in Dulbecco’s modified Eagle’s medium (DMEM) (Thermo Fisher Scientific) at 37°C in the atmosphere with 5% CO2. The retain KSHV genome, the iSLK/Bac16 cells were grown under selection in media with the addition of 150 μg/ml of hygromycin B, 1 μg/ml of puromycin and 250 μg/ml of G418. The lytic infection in iSLK/Bac16 cells was induced by adding 1 mM sodium butyrate (Millipore Sigma) and 1 μg/ml doxycycline (Thermo Fisher Scientific) for 48 h. Primary effusion lymphoma BCBL-1 cells carrying the KSHV genome were cultivated in RPMI 1640 medium (Thermo Fisher Scientific) at 37°C in the atmosphere with 5% CO2. KSHV lytic infection in BCBL-1 cells was induced by valproic acid (Sigma-Aldrich) treatment at a final concentration of 1 mM for 24 h. All media were supplemented with 10% fetal bovine serum (Cytiva) and 1 × penicillin-streptomycin-glutamine (Thermo Fisher Scientific). Both BCBL-1 and HEK293T cells were authenticated with a short tandem repeat (STR) profiling analysis performed by ATCC custom cell authentication services.

### RNA-seq and transcriptome analyses

The analysis of ORF57-associated RNAs (CLIP-seq) and differential expression of host genes in BCBL-1 with KSHV lytic versus latent infection were described in detail (46) and the associated sequencing data have been deposited in NCBI’s Gene Expression Omnibus (NCBI GEO) with the accession numbers (Acc. No.) GSE179726 and GSE179727.

To identify the differential expression of HEK293T cells with ORF57 expression versus cells transfected with the empty control vector, we performed mRNA-seq (NCBI GEO Acc. No. GSE179728). The obtained sequence reads in fastq format were mapped to the human reference genome (GRCh37) using STAR 2.7.6a (86). RSEM 1.3.0 (87) was used to quantify gene-level expression, with counts normalized to library size as counts-per-million. Limma-voom (88) was used for normalization and differential gene expression. Genes with a p-value ≤0.001, an adjustment of p-values by Benjamini Hochberg FDR (FDR < 0.05), and fold change values FC ≥ 2 for upregulated or FC≤2 for downregulated were considered statistically significant differentially expressed. The transcripts with normalized expression level reads per kilobase per million (RPKM) ≥3 were extracted for further analysis. Instant Clue software (89) and Heatmapper (90) online software (http://www.heatmapper.ca/) were used to generate the volcano plots and heatmaps, respectively.

To identify transcription factors driving detected transcriptome changes, we performed the genes set enrichment analysis (GSEA 4.1.0) using the gene ranking and TFT: transcription factors targets from C3:regulatory target gene sets (https://www.gsea-msigdb.org/).

### Reagents and antibodies

Specific reagents used in this study were actinomycin D, G418, hygromycin B, puromycin, sodium butyrate, TriPure Isolation Reagent solution, and valproic acid from Millipore Sigma, doxycycline and NuPAGE™ LDS Sample Buffer (4×) from Thermo Fisher Scientific, radioimmunoprecipitation assay buffer (RIPA) from Boston Bioproducts (Ashland, MA), and LipoD293™ In Vitro DNA Transfection Reagent from SignaGen Laboratories (Frederick, MD). Following antibodies were used in this report: affinity-purified rabbit polyclonal anti-ORF57 N-terminal (in house), mouse monoclonal anti-ORF57 N-terminal, and affinity-purified rabbit polyclonal anti-ORF57 C-terminal antibodies were from Rockland Immunochemicals. Rabbit polyclonal anti-AKT, rabbit monoclonal anti-p-AKT, rabbit monoclonal anti-c-FOS (9F6), mouse monoclonal anti-GAPDH, and rabbit polyclonal anti-c-JUN antibodies were purchased from Cell Signaling Technology. Rabbit polyclonal anti-RTA was a gift of Dr. Yoshi Izumiya (UC Davis). Mouse monoclonal anti-myc (9E10) was a gift of Dr. Xuefeng Liu (the Ohio State University). Mouse polyclonal anti-AEN was from Millipore Sigma, rabbit polyclonal anti-RGS2 from Abcam, rabbit polyclonal anti-ZFC3H1(MTR4) antibody from Novus biologicals, and rabbit IgG isotype, mouse IgG isotype were from Thermo Fisher Scientific.

### RNA immunoprecipitation (RIP) and RT-PCR

RIP was performed as described in previous reports for ORF57-CLIP-seq (46, 54). Cell lysates were prepared after washing cells twice with phosphate-buffered saline (PBS) by cell lysis in 1× RIPA buffer supplemented with a complete mini-EDTA-free protease inhibitor cocktail (Millipore Sigma) for 30 min on ice followed by brief sonication (10 short pulses, 1–2 seconds each, using a sonicator with a power setting at level 4). Lysates were then clarified by centrifugation at 20,000 × g for 15 min at 4 °C and pre-cleared by incubation with protein A-agarose beads coated with rabbit IgG for 2 h at 4°C. Protein A- agarose beads (Millipore Sigma) were washed 3 times with 1 × immunoprecipitation (IP) buffer (50 mM HEPES [pH 7.5], 200 mM NaCl, 1 mM EDTA, 2.5 mM EGTA, 10% glycerol, 0.1% NP-40) were coated with highly specific in house generated anti-ORF57 rabbit polyclonal antibody, anti-myc antibody or rabbit or mouse IgG isotype (Thermo Fisher Scientific), last two used as a negative control. The pre-cleared lysates were incubated with the antibody-coated beads overnight at 4°C, followed by extensive washes with 1 × IP buffer. Proteinase K (Millipore Sigma) was added to remove the proteins from RNA. The released RNA was extracted by phenol:chloroform and precipitated with sodium acetate. Total RNA from input cells was extracted by TRIpure reagent (Millipore Sigma). The cDNA synthesis was carried out by the SuperScript First-Strand Synthesis System kit (Thermo Fisher Scientific) in the absence (RT-) or presence (RT+) of reverse transcriptase (Applied Biosystems). The custom oligos for FOS (oBAH59 plus oBAH60 or oBAH55), PMF1(oBAH239 plus oBAH240), LY9 (oBAH233 plus oBAH234), and RBL1 (oBAH245 plus oBAH246) were used for mRNA detection by PCR with AmpliTaq DNA polymerase (Invitrogen).

### Cell extract preparation and RNA and DNA oligo pulldown assays

Cell extracts for oligo pulldown assays were isolated from HEK293T cells transfected with an ORF57-expressing vector (pVM7) using LipoD293 transfection reagent (SignaGen Laboratories). Twenty-four hours after transfection, the cells were washed with PBS and resuspended in ice-cold 1 × RIPA buffer containing protease inhibitors (Millipore Sigma). After brief sonication (10 short pulses, 1–2 seconds each, using a sonicator with a power setting at level 4), the cell extracts were cleared by 15 min centrifugation at 12,000 × g at 4 °C.

RNA oligo pulldown assays were performed using biotinylated RNA oligos (Integrated DNA Technologies, IDT) as previously described (46), with oBAH108, oBAH109, oBAH110, oBAH111, and oBAH156 derived from an ORF57 BS in the 5ʹ end (nt 179-237) of FOS mRNA (Genbank Acc. No. NM_005252.4). Previously reported oNP41 and oNP42 (29) derived from KSHV viral interleukin-6 (vIL6) RNA were used as a negative and positive control, respectively (see oligos table for details). 400 pmol of individual RNA oligos were immobilized on NeutrAvidin beads (Thermo Fisher Scientific) in 300 μl of 1 × Tris-Buffered Saline (TBS) at 4°C for 2 h. After two washes in 1 × TBS buffer, the oligo-coated beads were incubated with 100 μl cell extract obtained from ∼2 ×10^6^ cells. The reaction volume was adjusted to 400 μl with 1 × TBS buffer. After overnight incubation at 4°C on a rotating mixer, the beads were washed 3 times with 1 × TBS, and the proteins pulled down were eluted in 40 μl of 2 × LDS sample buffer (Thermo Fischer Scientific) supplemented with 5% 2-mercaptoethanol (2-ME). ORF57 in the pulldown was detected by Western blot with a rabbit polyclonal anti-ORF57 antibody (23).

Similarly, DNA pulldowns were performed using biotinylated double-stranded (ds) DNA oligos. To generate dsDNA oligos, equal amount of 200 µM of single-stranded complementary oligos was annealed by heating at 95 °C for 10 min followed by gradual cooling to room temperature. Following DNA oligos were used to generate DNA duplexes with a consensus AP-1-binding site (oBAH160 and oBAH161) and SP1-binding site (oHBL56 and oHBL57) or FOS-binding site A (oBAH162 and oBAH163), FOS-binding site B (oBAH167 and oBAH168), and FOS-binding site C (oBAH169 and oBAH170) from the *RSG2* promoter. Before pulldown, 100 µl (50% slurry) of Pierce NeutrAvidin Agarose beads (Thermo Fisher Scientific) were washed 3 times and re-suspended in 500 ml of 1 × wash buffer (20 mM Tris-HCl [pH 7.5], 100 mM NaCl, 1 mM MgCl2, 0.5 mM EDTA, 0.5 mM DTT) and incubated with 10 µg of biotinylated dsDNA oligos for 2 h at room temperature. After 3 washes with the wash buffer, the oligos-coated beads were mixed with total cell lysate from ∼2 x 10^6^ cells in 500 µl of pulldown buffer (20 mM Tris-HCl [pH7.5], 100 mM NaCl, 1 mM MgCl2, 0.5 mM EDTA, 0.5 mM DTT, 4% glycerol, 10 µg/ml of poly dI-dC [Sigma], 1 × protease inhibitor cocktail [Millipore Sigma]) and incubated overnight at 4°C. The unbound proteins were removed by an extensive wash with the wash buffer. Finally, the proteins pulled down were eluted with 50 μl of 2 × LDS protein sample buffer supplemented with 2-ME (5%) and resolved by SDS-PAGE for Western blot analysis with the anti-c-FOS (9F6) or anti-c-JUN rabbit polyclonal antibodies.

### RT-qPCR

Total cell RNA from BCBL-1, HEK293T and iSLKBac16 was extracted using TRIpure reagent (Millipore Sigma) following company instructions. cDNA synthesis was carried out with SuperScript First-Strand Synthesis System kit (Thermo Fisher Scientific). qPCR was performed with TaqMan Gene Expression Assays (Thermo Fisher Scientific) for FOS, RBL1, RGS2, AEN, MTR4, and GADPH in TaqMan Gene Expression Master Mix (Applied Biosystems) at StepOne Plus Real-Time PCR system (Applied Biosystems). The changes in transcript abundance were analyzed using the 2−ΔΔCt method and normalized to GAPDH levels. All samples were analyzed in triplicates.

### FOS RNA transcriptional pulse-chase assay

HEK293T cells (5 × 10^5^) seed in a 6-well plate were transiently co-transfected with 200 ng of a myc-tagged FOS (pCMV6-FOS, OriGene, RC202597) and 600 ng of a wt ORF57 (pVM7) expression vector or an empty FLAG control vector (pFLAG-CMV-5.1, Millipore Sigma). After 24 h of transfection, the transcription was stopped by adding 15 μg/ml (final concentration) of RNA polymerase II (Pol II) inhibitor actinomycin D (Act. D). The total RNA samples were collected at 0.5, 1, 3, 6 and 9 h after Act. D addition usingTriPure reagent. Northern blot analysis using 5 μg of total RNA of each sample was performed as previously described (22). The ^32^P end-labeled antisense oXL39 and oZMZ270 were used to detect FOS-myc and GAPDH mRNA, respectively. The FOS-myc RNA decay curve was constructed based on a densitometry quantification of individual FOS RNA band in each sample after GAPDH normalization with the value at t = 0 (Act. D addition time) setting as 100%. The decay curves were then fitted to an exponential decay model to calculate the half-time of the corresponding transcripts.

### Northern blot for detection of endogenous or exogenous FOS RNA

Total RNA (8 μg) in each group extracted from HEK293 cells with or without AEN knockdown or with or without Myc-tagged FOS expression was separated in a 1% (wt/vol) formaldehyde-agarose gel. The gel was transferred to a nylon membrane and immobilized by UV light. The membrane was prehybridized 1 h and followed by 24 h hybridization at 42°C using a ^32^P labeled antisense oligo oBAH55 for detection of endogenous FOS and oZMZ270 for GAPDH serving as a sample loading control.

### Expression vectors, promoter-reporter plasmids, and dual-luciferase assays

Myc-tagged FOS (pCMV6-FOS) expressing vector was purchased from OriGene. The pBAH4 plasmid with point mutations in the ORF57 BS in FOS cDNA was generated by overlapping PCR using a set of primers with mutated BS (oBAH138 and oBAH139) in combination with flanking primers oBAH137 and oBAH141 for individual PCRs. The amplified product of the expected size was then back cloned into the pCMV6-FOS-Myc by using *Bam*HI and *Apa*I restriction sites.

To evaluate the activity of the *RGS2* promoter, a 1.5-kb long DNA fragment covering −1400 to +100 nt from the annotated TSS corresponding to a genomic region of chr1:192776769-192778268 (GRCh37/hg19) was amplified by PCR from HEK293T cell genomic DNA using the oligos oBAH173 and oBAH174 and cloned into a Firefly luciferase reporter, pGL3-basic vector (Promega) via *Sma*I and *Hind*III sites to generate pBAH8. The overlapping PCR with mutated oBAH184 and oBAH186 oligos combined with oBAH173 and oBAH183 were used to generate plasmid pBAH9 containing the *RGS2* promoter with mutated FOS-binding motif A. Each plasmid was verified by restriction enzyme digestion and DNA Sanger sequencing.

For dual-luciferase assays, 1 × 10^5^ of HEK293T cells seeded in a 24-well plate were co-transfected with individual Firefly luciferase reporter plasmid together with a Renilla luciferase plasmid, pRL-TS (36, 91), using LipoD293 DNA transfection reagent (SignaGen) in triplicates. Dual-luciferase reporter assays (Promega) were performed according to the manufacturer’s instructions. Each sample’s relative Firefly luciferase activity was normalized to Renilla luciferase (Rluc) activity.

### siRNA-mediated knockdown

To knock down the endogenous expression of host genes or ectopic viral ORF57 expression in HEK293T cells, 5 ×10^5^ cells seeded in a 6-well plate were transfected with 40 nM SMARTpool human siRNAs (Horizon) or non-targeting (siNT) control siRNA (Horizon) using LipoJet In Vitro Transfection Kit (Ver. II) (SignaGen Laboratories). Twenty-four hours after the first transfection, the cells were divided into two wells and subjected to the second round of siRNA transfection, followed by transfection of ORF57- expression or empty vectors. After 48 h, total RNA extraction was extracted by TriPure reagent for RT-qPCR. Protein lysates obtained by direct cell lysis in 2 × LDS protein sample buffer supplemented with 5% (vol/vol) 2-ME were analyzed by Western blotting for the protein expression of FOS, MTR4, RGS2, AKT, pAKT, GAPDH, and ORF57.

### Co-Immunoprecipitation assay

HEK293T (1 × 10^7^) cells transfected with 8 µg of ORF57-expressing vector (pVM7), AEN- Myc, or a mixture of AEN-myc andORF57 (1:1) by using LipoD293. After 24 h of transfection, the cells were washed once with cold PBS and lysed in 500 μl of ice-cold 1 × RIPA buffer. The obtained protein lysates were pre-cleared by centrifugation and incubated with 100 μl (50% slurry) of protein A beads (Millipore Sigma), coated specific antibodies or corresponding species-specific IgG isotype control in 1 ml of 1 × IP buffer. The mixture was incubated overnight at 4°C followed by 3 washes with IP buffer. The immunoprecipitated complexes were eluted with 2 × LDS protein sample buffer supplemented with 10% (vol/vol) 2-ME and analyzed by Western blotting to detect individual cellular or viral proteins using specific antibodies.

### In vitro transcription and translation assays

In vitro transcription and translation reactions using rabbit reticulocyte lysates as previously described (29) were performed by TNT Quick Coupled Transcription/Translation System (Promega) in the presence of [^35^S] Met (Perkin-Elmer). *FOS* cDNA under the T7 promoter was used as a template (OriGene). Following manufacturer’s instructions, the Firefly luciferase RNA was included as a translation control responder. The purified baculovirus-expressed recombinant FLAG-tagged ORF57 protein (21) was added to the translational reaction at increasing concentrations (5-50 nM). The translation was performed for 90 min at 30°C and stopped by the addition of 2 × LDS sample buffer and heat inactivation at 80 °C for 10 min. Translated proteins labeled with [^35^S] Met were resolved in a 4%-12% SDS-PAGE gel (Invitrogen), dried onto a Whatman filter paper, and exposed to a phosphor screen or X-ray film for quantification.

### Lentivirus production and transduction

HEK293T cells, grown to ∼70% confluency, were co-transfected with ORF57 cDNA cloned in pLenti-P2A-Puro Lentiviral Gene Expression Vector (OriGene) together with pCMV.Dr8.2 and pCMV.VSV.G in a ratio of 10:10:3.5 using Lipofectamine 2000 (ThermoFisher Scientific). Lentiviral supernatant was harvested 48 h later and then filtered to be concentrated through a Lenti-X Concentrator (Takara) following manufacturer’s instructions. The concentrated virus preps were resuspended in 1 × PBS and titrated by quantification of the p24 capsid protein using a standard p24 antigen ELISA kit. About 2.5 × 10^6^ of the 57KO iSLK/Bac16 cells were seeded in a 6-well plate and infected with the ORF57 lentiviral particles at 10 MOI (multiplicity of infection) in the presence of 8 μg/mL polybrene. After 4 h of infection, the infectious virus particles were removed by rinsing the cells with 1 × PBS and replaced with fresh culture media. Lytic KSHV infection in the 57KO iSLK/Bac16 cells was induced by butyrate and doxycycline for 48 h. Total cell RNA was extracted with TRIpure reagent and total protein lysate was prepared by adding 150 ul of 2 × LDS supplemented with 5% of 2-ME.

### Statistics

All statistical analyses were performed by unpaired, two-tailed Student t-test, with *, p <0.05; **, p < 0.01; ***, p < 0.001, NS – no-significance.

### Data, Materials, and Software Availability

The RNA-seq data generated in this study are available at the NCBI’s Gene Expression Omnibus (NCBI GEO) with the accession numbers (Acc. No.) GSE179726, GSE179727, and GSE179728. All unique reagents generated during this study are available from the Lead contact without restriction.

## SUPPLEMENTAL INFORMATION

Table S1. KSHV ORF57 host targets in BCBL-1 cells identified by ORF57 CLIP-seq (this study) and ORF57 HITS-CLIP (Sei et al., 2015, PLoS Pathog 11(2): e1004652; doi.org/10.1371/journal.ppat.1004652).

Table S2. Peaks representing ORF57-binding sites in transcripts of host protein-coding genes identified by Piranha software based on anti-ORF57 CLIP-seq as shown in Fig. 1C. The associated p-value represents zero-truncated negative binomial distribution. The peaks with a p-value less than 0.05 are considered significant.

Table S3. Differential expression of host coding and noncoding genes in BCBL-1 cells with KSHV latent versus lytic infection.

Table S4. Differential expression of coding and noncoding genes in HEK293T cells without versus with KSHV ORF57.

Table S5. AP-1 sensitive genes identified by GSEA from BCBL-1 cells with KSHV lytic infection.

Table S6. Oligonucleotides used in the study.

## Supplemental Figures

**Figure S1.**
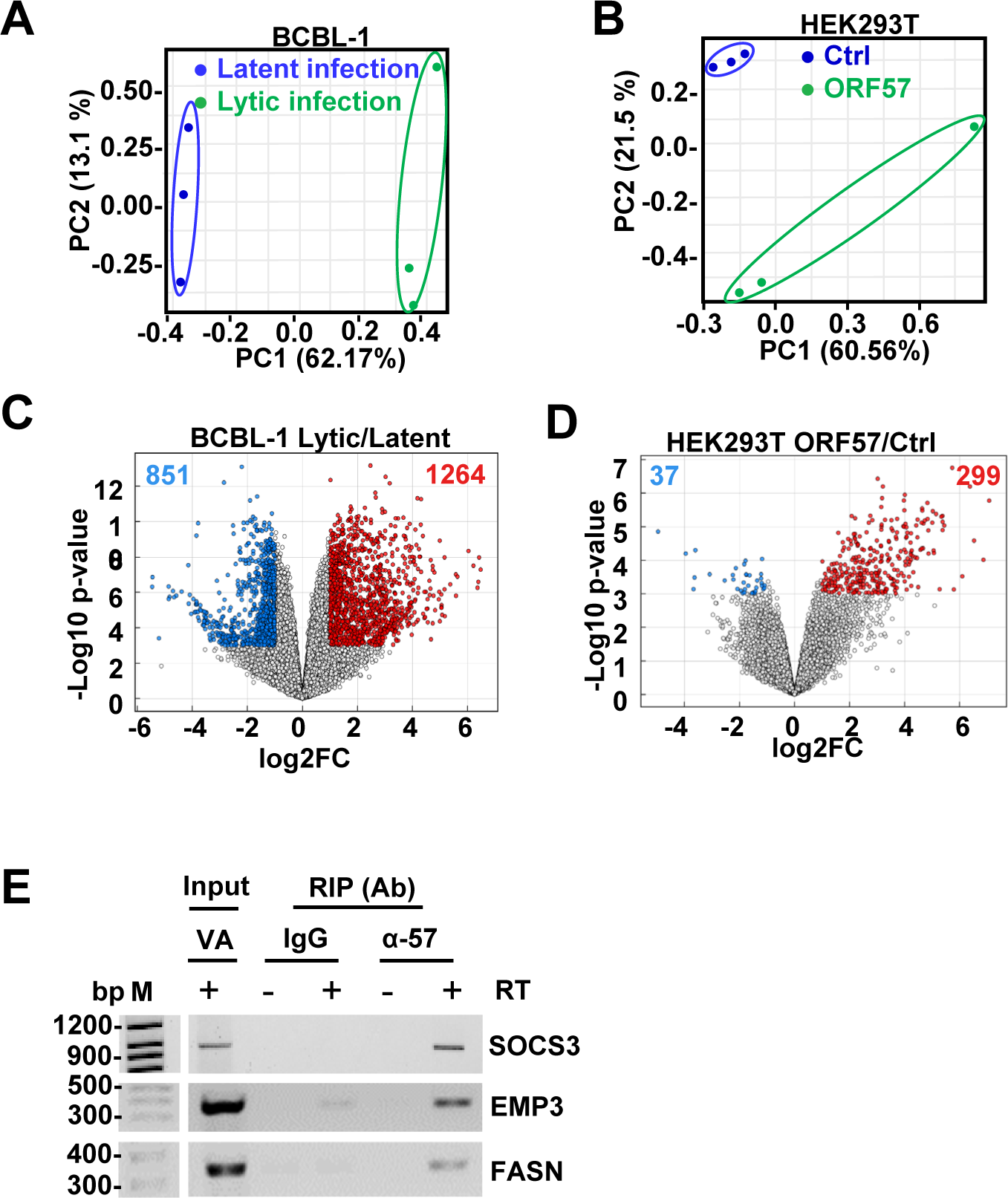
ORF57 regulates the expression of host protein-coding RNAs. (A and B) Principal component (PC) analysis of RNA-seq of BCBL-1 cells with latent or lytic infection (A) and HEK293T cells without (ctrl) or with ORF57 expression (B); each point represents individual samples in each group. (C and D) Volcano plots of differentially expressed host genes in BCBL-1 cells with lytic versus latent infection (C) and HEK293T cells with ORF57 expression over the control cells without ORF57 (D). Genes significantly (p<0.001) upregulated in FC (fold change ≥ 2) are shown as red dots and downregulated FC ≤ −2 as blue dots. (E) Validation of interaction between ORF57 and SOCS3, EMP3, and FASN RNA by RIP. RT-PCR in the presence (+) or absence (-) of RT with a gene-specific primer set (Table 1) was performed on RNA immunoprecipitated (RIP) from BCBL-1 cells during KSHV lytic infection by anti-ORF57 antibody. Corresponding non-specific IgG served as a negative control.

**Figure S2.**
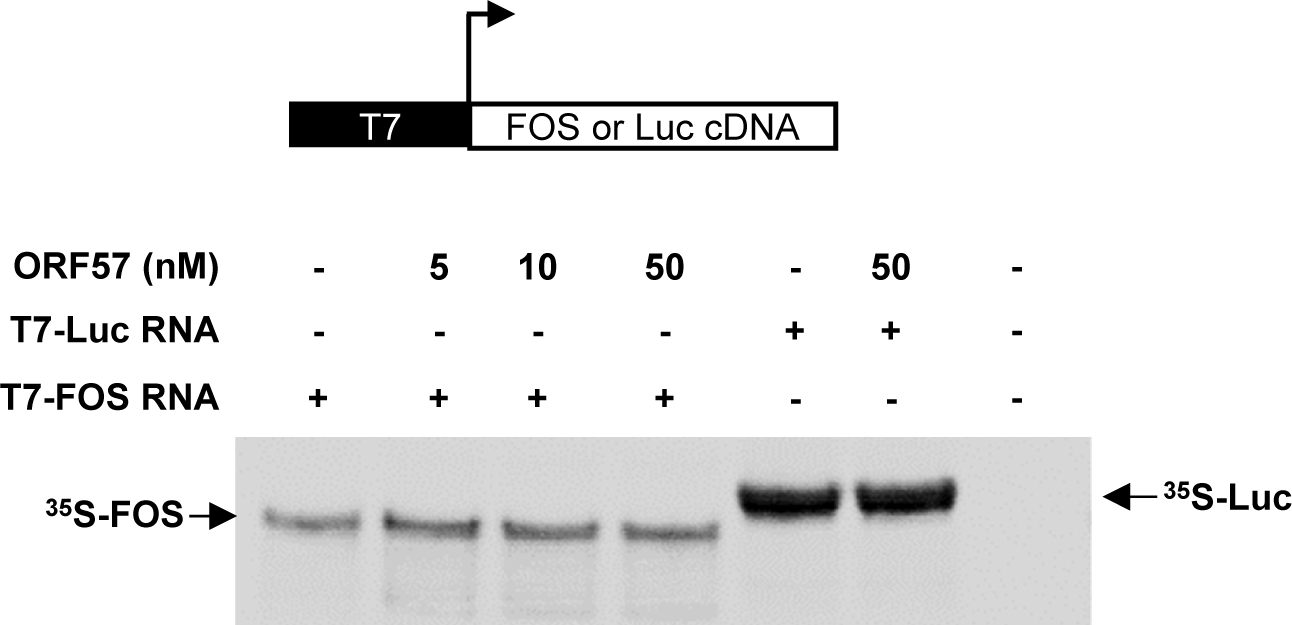
FOS expression is not regulated by ORF57 at the translation level. A) A representative gel of two in vitro transcription/translation assays using rabbit reticulocyte lysates in the presence of ^35^S-methionine. *FOS* cDNA under the T7 promoter was used as a template. The recombinant FLAG-tagged ORF57 protein was added at increasing concentrations (5-50 nM) in the reaction and incubated at 30 °C for 1.5 h. The reaction was stopped by the addition of 2 × LDS/2-ME protein sample buffer and heated at 80 °C for 10 min. ^35^S-met-labeled proteins were resolved in SDS-PAGE gel and captured by phosphorimager screen or X-ray film. Firefly luciferase RNA (Luc) was included as a negative control.

**Figure S3.**
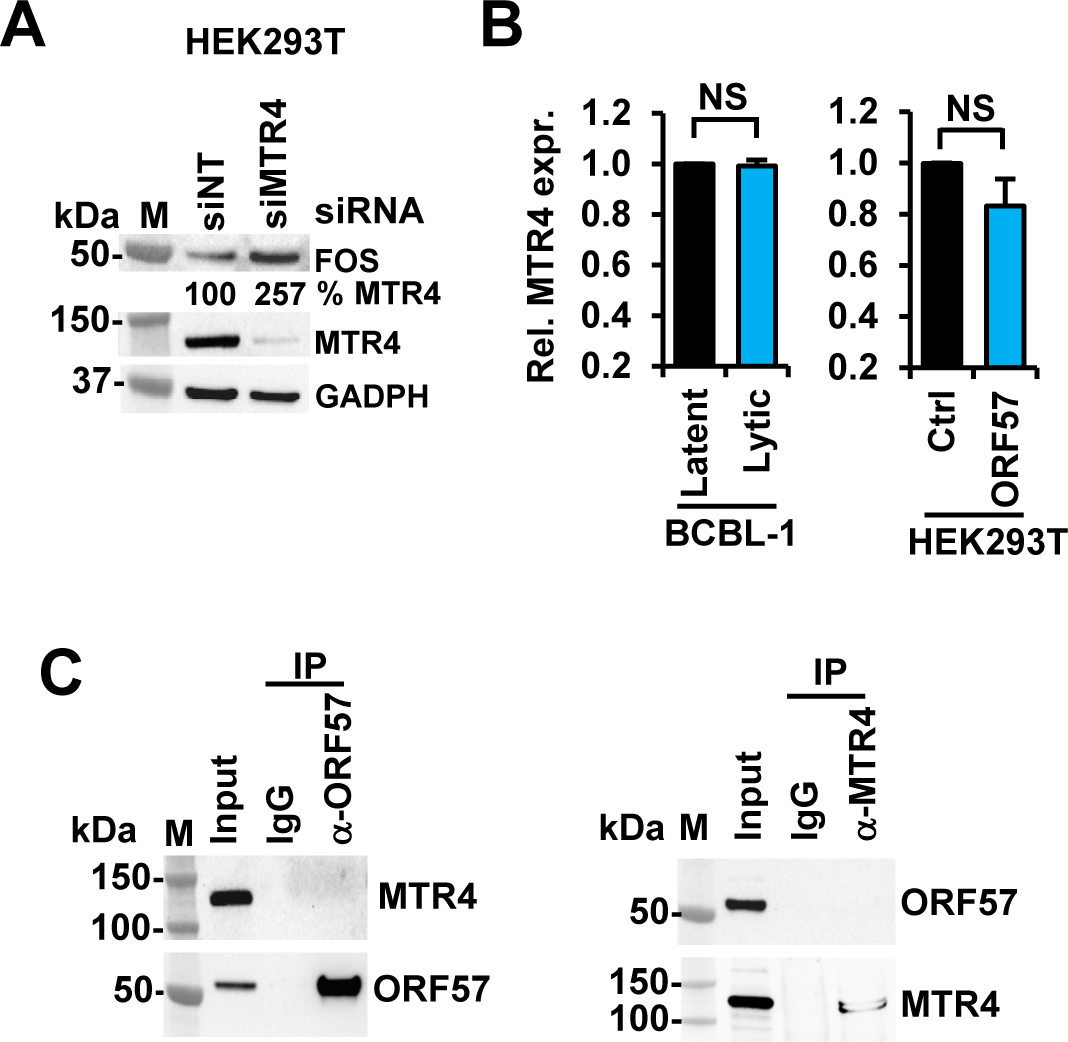
FOS expression is sensitive to host MTR4. (A) MTR4 knockdown (KD) by MTR4-targeting siRNA (siMTR4) promotes FOS protein expression in HEK293T cells. The effect of MTR4 KD on FOS expression in HEK293T cells was examined by Western blotting at 48 h after transfection. A non-targeting siRNA (siNT) served as a siRNA control and GAPDH as a sample loading control. (B) KSHV lytic infection and ORF57 expression do not affect MTR4 expression. MTR4 RNA expression was quantified by RT-qPCR in BCBL-1 with latent and lytic infection and in HEK293T cells transfected with an ORF57-expressing (ORF57) or an empty vector control (Ctrl) vector. Data from all three separate experiments, each with three replicates, were averaged with mean ± SD. NS – no significance, two-tailed Student *t*-test. (C) No protein-protein interaction between ORF57 and MTR4. Co-immunoprecipitation of ORF57 and MTR4 proteins from total cell extracts from HEK293T cells transfected with an ORF57-expressing vector was performed by using rabbit anti-ORF57 or anti-MTR4 antibodies, with corresponding species-specific IgG isotype serving as an antibody control. ORF57 and MTR4 proteins were immunoblotted with the corresponding antibodies as indicated.

## REFERENCES

1. D. Ganem, KSHV and the pathogenesis of Kaposi sarcoma: listening to human biology and medicine. J. Clin. Invest 120, 939–949 (2010).

2. E. A. Mesri, E. Cesarman, C. Boshoff, Kaposi’s sarcoma and its associated herpesvirus. Nat. Rev. Cancer 10, 707–719 (2010).

3. Y. Chang et al., Identification of herpesvirus-like DNA sequences in AIDS-associated Kaposi’s sarcoma. Science 266, 1865–1869 (1994).

4. C. Vogt, J. Bohne, The KSHV RNA regulator ORF57: target specificity and its role in the viral life cycle. Wiley Interdiscip Rev RNA 7, 173–185 (2016).

5. E. Shaulian, M. Karin, AP-1 in cell proliferation and survival. Oncogene 20, 2390–2400 (2001).

6. R. Eferl, E. F. Wagner, AP-1: a double-edged sword in tumorigenesis. Nat Rev Cancer 3, 859–868 (2003).

7. N. Szaloki, J. W. Krieger, I. Komaromi, K. Toth, G. Vamosi, Erratum for Szaloki et al., “Evidence for Homodimerization of the c-Fos Transcription Factor in Live Cells Revealed by Fluorescence Microscopy and Computer Modeling”. Mol Cell Biol 38, e00282 (2018).

8. H. Gazon, B. Barbeau, J. M. Mesnard, J. M. Peloponese, Jr., Hijacking of the AP- 1 Signaling Pathway during Development of ATL. Front Microbiol 8, 2686 (2017).

9. H. Mirzaei et al., Histone deacetylases in virus-associated cancers. Rev Med Virol 30, e2085 (2020).

10. S. E. Wang et al., Early activation of the Kaposi’s sarcoma-associated herpesvirus RTA, RAP, and MTA promoters by the tetradecanoyl phorbol acetate-induced AP1 pathway. J. Virol 78, 4248-4267 (2004).

11. C. Alfonso-Gonzalez, J. R. Riesgo-Escovar, Fos metamorphoses: Lessons from mutants in model organisms. Mech Dev 154, 73–81 (2018).

12. J. Xie, H. Pan, S. Yoo, S. J. Gao, Kaposi’s sarcoma-associated herpesvirus induction of AP-1 and interleukin 6 during primary infection mediated by multiple mitogen-activated protein kinase pathways. J Virol 79, 15027–15037 (2005).

13. N. Sharma-Walia et al., ERK1/2 and MEK1/2 induced by Kaposi’s sarcoma-associated herpesvirus (human herpesvirus 8) early during infection of target cells are essential for expression of viral genes and for establishment of infection. J Virol 79, 10308–10329 (2005).

14. H. Pan, J. Xie, F. Ye, S. J. Gao, Modulation of Kaposi’s sarcoma-associated herpesvirus infection and replication by MEK/ERK, JNK, and p38 multiple mitogen-activated protein kinase pathways during primary infection. J Virol 80, 5371-5382 (2006).

15. X. Li et al., ORF45-Mediated Prolonged c-Fos Accumulation Accelerates Viral Transcription during the Late Stage of Lytic Replication of Kaposi’s Sarcoma-Associated Herpesvirus. J Virol 89, 6895–6906 (2015).

16. V. Majerciak, Z. M. Zheng, KSHV ORF57, a Protein of Many Faces. Viruses 7, 604–633 (2015).

17. A. BeltCappellino, et al., CRISPR/Cas9-mediated knockout and in situ inversion of ORF57 gene from all copies of the KSHV genome in BCBL-1 cells. J. Virol 93, e00628 (2019).

18. O. Manners, J. C. Murphy, A. Coleman, D. J. Hughes, A. Whitehouse, Contribution of the KSHV and EBV lytic cycles to tumourigenesis. Curr Opin Virol 32, 60–70 (2018).

19. J. R. Kirshner, D. M. Lukac, J. Chang, D. Ganem, Kaposi’s sarcoma-associated herpesvirus open reading frame 57 encodes a posttranscriptional regulator with multiple distinct activities. J. Virol 74, 3586–3597 (2000).

20. A. K. Gupta, V. Ruvolo, C. Patterson, S. Swaminathan, The human herpesvirus 8 homolog of Epstein-Barr virus SM protein (KS-SM) is a posttranscriptional activator of gene expression. J. Virol 74, 1038–1044 (2000).

21. V. Majerciak, M. Kruhlak, P. K. Dagur, J. P. McCoy, Jr., Z. M. Zheng, Caspase-7 cleavage of Kaposi sarcoma-associated herpesvirus ORF57 confers a cellular function against viral lytic gene expression. J. Biol. Chem 285, 11297–11307 (2010).

22. V. Majerciak, K. Yamanegi, S. H. Nie, Z. M. Zheng, Structural and functional analyses of Kaposi sarcoma-associated herpesvirus ORF57 nuclear localization signals in living cells. J. Biol. Chem 281, 28365–28378 (2006).

23. V. Majerciak, N. Pripuzova, J. P. McCoy, S. J. Gao, Z. M. Zheng, Targeted disruption of Kaposi’s sarcoma-associated herpesvirus ORF57 in the viral genome is detrimental for the expression of ORF59, K8alpha, and K8.1 and the production of infectious virus. J. Virol 81, 1062–1071 (2007).

24. Z. Han, S. Swaminathan, Kaposi’s sarcoma-associated herpesvirus lytic gene ORF57 is essential for infectious virion production. J. Virol 80, 5251–5260 (2006).

25. V. Majerciak, K. Yamanegi, Z. M. Zheng, Gene structure and expression of Kaposi’s sarcoma-associated herpesvirus ORF56, ORF57, ORF58, and ORF59. J. Virol 80, 11968–11981 (2006).

26. F. Yuan et al., The crystal structure of KSHV ORF57 reveals dimeric active sites important for protein stability and function. PLoS Pathog 14, e1007232 (2018).

27. K. Nishimura et al., A posttranscriptional regulator of Kaposi’s sarcoma-associated herpesvirus interacts with RNA-binding protein PCBP1 and controls gene expression through the IRES. Virology 325, 364–378 (2004).

28. V. Majerciak et al., Kaposi’s sarcoma-associated herpesvirus ORF57 interacts with cellular RNA export cofactors RBM15 and OTT3 to promote expression of viral ORF59. J. Virol 85, 1528–1540 (2011).

29. J. G. Kang et al., Kaposi’s Sarcoma-Associated Herpesvirus ORF57 Promotes Escape of Viral and Human Interleukin-6 from MicroRNA-Mediated Suppression. J. Virol 85, 2620–2630 (2011).

30. N. R. Sharma et al., KSHV RNA-binding protein ORF57 inhibits P-body formation by interaction with Ago2 and GW182. Nucleic Acids Res 47, 9368–9385 (2019).

31. N. R. Sharma, V. Majerciak, M. J. Kruhlak, Z. M. Zheng, KSHV inhibits stress granule formation by viral ORF57 blocking PKR activation. PLoS Pathog 13, e1006677 (2017).

32. M. Nekorchuk, Z. Han, T. T. Hsieh, S. Swaminathan, Kaposi’s Sarcoma-Associated Herpesvirus ORF57 Protein Enhances mRNA Accumulation Independently of Effects on Nuclear RNA Export. J. Virol 81, 9990–9998 (2007).

33. D. J. Li, D. Verma, S. Swaminathan, Binding of cellular export factor REF/Aly by Kaposi’s sarcoma-associated herpesvirus (KSHV) ORF57 protein is not required for efficient KSHV lytic replication. J. Virol 86, 9866–9874 (2012).

34. B. B. Sahin, D. Patel, N. K. Conrad, Kaposi’s sarcoma-associated herpesvirus ORF57 protein binds and protects a nuclear noncoding RNA from cellular RNA decay pathways. PLoS. Pathog 6, e1000799 (2010).

35. G. R. Pilkington et al., Kaposi’s Sarcoma-Associated Herpesvirus ORF57 Is Not a Bona Fide Export Factor. J. Virol 86, 13089–13094 (2012).

36. M. J. Massimelli et al., Stability of a Long Noncoding Viral RNA Depends on a 9- nt Core Element at the RNA 5’ End to Interact with Viral ORF57 and Cellular PABPC1. Int. J. Biol. Sci 7, 1145–1160 (2011).

37. M. J. Massimelli, V. Majerciak, M. Kruhlak, Z. M. Zheng, Interplay between polyadenylate-binding protein 1 and Kaposi’s sarcoma-associated herpesvirus ORF57 in accumulation of polyadenylated nuclear RNA, a viral long noncoding RNA. J. Virol 87, 243–256 (2013).

38. M. J. Massimelli et al., Multiple Regions of Kaposi’s Sarcoma-Associated Herpesvirus ORF59 RNA are Required for Its Expression Mediated by Viral ORF57 and Cellular RBM15. Viruses 7, 496–510 (2015).

39. E. Sei, N. K. Conrad, Delineation of a core RNA element required for Kaposi’s sarcoma-associated herpesvirus ORF57 binding and activity. Virology 419, 107–116 (2011).

40. V. Majerciak et al., Kaposi’s sarcoma-associated herpesvirus ORF57 functions as a viral splicing factor and promotes expression of intron-containing viral lytic genes in spliceosome-mediated RNA splicing. J. Virol 82, 2792–2801 (2008).

41. V. Majerciak et al., Stability of structured Kaposi’s sarcoma-associated herpesvirus ORF57 protein is regulated by protein phosphorylation and homodimerization. J. Virol 89, 3256–3274 (2015).

42. V. Majerciak, M. Lu, X. Li, Z. M. Zheng, Attenuation of the suppressive activity of cellular splicing factor SRSF3 by Kaposi sarcoma-associated herpesvirus ORF57 protein is required for RNA splicing. RNA 20, 1747–1758 (2014).

43. J. G. Kang et al., Kaposi’s sarcoma-associated herpesviral IL-6 and human IL-6 open reading frames contain miRNA binding sites and are subject to cellular miRNA regulation. J. Pathol 225, 378–389 (2011).

44. E. Sei, T. Wang, O. V. Hunter, Y. Xie, N. K. Conrad, HITS-CLIP analysis uncovers a link between the Kaposi’s sarcoma-associated herpesvirus ORF57 protein and host pre-mRNA metabolism. PLoS. Pathog 11, e1004652 (2015).

45. K. L. Harper et al., Dysregulation of the miR-30c/DLL4 axis by circHIPK3 is essential for KSHV lytic replication. EMBO Rep 23, e54117 (2022).

46. B. Alvarado-Hernandez et al., Protein-RNA Interactome Analysis Reveals Wide Association of Kaposi’s Sarcoma-Associated Herpesvirus ORF57 with Host Noncoding RNAs and Polysomes. J Virol 96, e0178221 (2022).

47. B. R. Jackson, M. Noerenberg, A. Whitehouse, A novel mechanism inducing genome instability in Kaposi’s sarcoma-associated herpesvirus infected cells. PLoS. Pathog 10, e1004098 (2014).

48. D. Palmeri, S. Spadavecchia, K. D. Carroll, D. M. Lukac, Promoter- and cell-specific transcriptional transactivation by the Kaposi’s sarcoma-associated herpesvirus ORF57/Mta protein. J. Virol 81, 13299–13314 (2007).

49. K. G. Eby et al., ISG20L1 is a p53 family target gene that modulates genotoxic stress-induced autophagy. Mol Cancer 9, 95 (2010).

50. S. Zou et al., ISG20L1 acts as a co-activator of DAPK1 in the activation of the p53- dependent cell death pathway. J Cell Sci 136 (2023).

51. T. Kawase et al., p53 target gene AEN is a nuclear exonuclease required for p53- dependent apoptosis. Oncogene 27, 3797–3810 (2008).

52. A. Linder, M. Hagberg Thulin, J. E. Damber, K. Welen, Analysis of regulator of G- protein signalling 2 (RGS2) expression and function during prostate cancer progression. Sci Rep 8, 17259 (2018).

53. S. J. Lin et al., RGS2 Suppresses Melanoma Growth via Inhibiting MAPK and AKT Signaling Pathways. Anticancer Res 41, 6135–6145 (2021).

54. Y. Ma, P. Liu, V. Majerciak, J. Zhu, Z. M. Zheng, CLIP-seq to Identify KSHV ORF57-Binding RNA in Host B Cells. Curr Protoc Microbiol 41, 1E 11 11–11E 11 18 (2016).

55. P. J. Uren et al., Site identification in high-throughput RNA-protein interaction data. Bioinformatics 28, 3013–3020 (2012).

56. U. Raudvere et al., g:Profiler: a web server for functional enrichment analysis and conversions of gene lists (2019 update). Nucleic Acids Res 47, W191–W198 (2019).

57. Z. Yin et al., The essential role of Cited2, a negative regulator for HIF-1alpha, in heart development and neurulation. Proc Natl Acad Sci U S A 99, 10488–10493 (2002).

58. S. J. Freedman et al., Structural basis for negative regulation of hypoxia-inducible factor-1alpha by CITED2. Nat Struct Biol 10, 504–512 (2003).

59. J. A. Castro-Mondragon et al., JASPAR 2022: the 9th release of the open-access database of transcription factor binding profiles. Nucleic Acids Res 50, D165–D173 (2021).

60. R. Chiu, P. Angel, M. Karin, Jun-B differs in its biological properties from, and is a negative regulator of, c-Jun. Cell 59, 979–986 (1989).

61. J. L. Meier, M. F. Stinski, Regulation of human cytomegalovirus immediate-early gene expression. Intervirology 39, 331–342 (1996).

62. H. Isomura, M. F. Stinski, The human cytomegalovirus major immediate-early enhancer determines the efficiency of immediate-early gene transcription and viral replication in permissive cells at low multiplicity of infection. J Virol 77, 3602–3614 (2003).

63. J. C. Ruiz, O. V. Hunter, N. K. Conrad, Kaposi’s sarcoma-associated herpesvirus ORF57 protein protects viral transcripts from specific nuclear RNA decay pathways by preventing hMTR4 recruitment. PLoS Pathog 15, e1007596 (2019).

64. E. M. Weick et al., Helicase-Dependent RNA Decay Illuminated by a Cryo-EM Structure of a Human Nuclear RNA Exosome-MTR4 Complex. Cell 173, 1663–1677.e1621 (2018).

65. F. Mure et al., The splicing factor SRSF3 is functionally connected to the nuclear RNA exosome for intronless mRNA decay. Sci Rep 8, 12901 (2018).

66. M. Lingaraju et al., The MTR4 helicase recruits nuclear adaptors of the human RNA exosome using distinct arch-interacting motifs. Nature Communications 10, 3393 (2019).

67. A. Subramanian et al., Gene set enrichment analysis: a knowledge-based approach for interpreting genome-wide expression profiles. Proc Natl Acad Sci U S A 102, 15545–15550 (2005).

68. A. J. Kimple et al., Structural determinants of G-protein alpha subunit selectivity by regulator of G-protein signaling 2 (RGS2). J Biol Chem 284, 19402–19411 (2009).

69. D. Verma, D. J. Li, B. Krueger, R. Renne, S. Swaminathan, Identification of the Physiological Gene Targets of the Essential Lytic Replicative Kaposi’s Sarcoma-Associated Herpesvirus ORF57 Protein. J. Virol 89, 1688–1702 (2015).

70. X. Chen, S. A. Castro, Q. Liu, W. Hu, S. Zhang, Practical considerations on performing and analyzing CLIP-seq experiments to identify transcriptomic-wide RNA-protein interactions. Methods 155, 49–57 (2019).

71. F. C. Y. Lee, J. Ule, Advances in CLIP Technologies for Studies of Protein-RNA Interactions. Mol Cell 69, 354–369 (2018).

72. M. Hafner et al., CLIP and complementary methods. Nature Reviews Methods Primers 1, 20 (2021).

73. L. O. Murphy, S. Smith, R. H. Chen, D. C. Fingar, J. Blenis, Molecular interpretation of ERK signal duration by immediate early gene products. Nat Cell Biol 4, 556–564 (2002).

74. S. Hollinger, J. R. Hepler, Cellular regulation of RGS proteins: modulators and integrators of G protein signaling. Pharmacol Rev 54, 527–559 (2002).

75. H. J. McNabb, Q. Zhang, B. Sjogren, Emerging Roles for Regulator of G Protein Signaling 2 in (Patho)physiology. Mol Pharmacol 98, 751–760 (2020).

76. J. H. Kehrl, S. Sinnarajah, RGS2: a multifunctional regulator of G-protein signaling. Int J Biochem Cell Biol 34, 432–438 (2002).

77. P. Osei-Owusu, K. J. Blumer, Regulator of G Protein Signaling 2: A Versatile Regulator of Vascular Function. Prog Mol Biol Transl Sci 133, 77–92 (2015).

78. T. Ingi et al., Dynamic regulation of RGS2 suggests a novel mechanism in G- protein signaling and neuronal plasticity. J Neurosci 18, 7178–7188 (1998).

79. K. J. Perschbacher et al., Reduced mRNA Expression of RGS2 (Regulator of G Protein Signaling-2) in the Placenta Is Associated With Human Preeclampsia and Sufficient to Cause Features of the Disorder in Mice. Hypertension 75, 569–579 (2020).

80. J. Cho et al., RGS2-mediated translational control mediates cancer cell dormancy and tumor relapse. J Clin Invest 131, e136779 (2021).

81. C. H. Nguyen, P. Zhao, A. J. Sobiesiak, P. Chidiac, RGS2 is a component of the cellular stress response. Biochem Biophys Res Commun 426, 129–134 (2012).

82. C. H. Nguyen et al., Translational control by RGS2. J Cell Biol 186, 755–765 (2009).

83. C. J. Wang, P. Chidiac, RGS2 promotes the translation of stress-associated proteins ATF4 and CHOP via its eIF2B-inhibitory domain. Cell Signal 59, 163–170 (2019).

84. S. Tang, K. Yamanegi, Z. M. Zheng, Requirement of a 12-base-pair TATT-containing sequence and viral lytic DNA replication in activation of the Kaposi’s sarcoma-associated herpesvirus K8.1 late promoter. J. Virol 78, 2609–2614 (2004).

85. 85. K. F. Brulois, et al., Construction and manipulation of a new Kaposi’s sarcoma-associated herpesvirus bacterial artificial chromosome clone. *J. Virol* 86, 9708- 9720 (2012).

86. A. Dobin et al., STAR: ultrafast universal RNA-seq aligner. Bioinformatics 29, 15–21 (2013).

87. B. Li, C. N. Dewey, RSEM: accurate transcript quantification from RNA-Seq data with or without a reference genome. BMC Bioinformatics 12, 323 (2011).

88. C. W. Law, Y. Chen, W. Shi, G. K. Smyth, voom: Precision weights unlock linear model analysis tools for RNA-seq read counts. Genome Biol 15, R29 (2014).

89. H. Nolte, T. D. MacVicar, F. Tellkamp, M. Kruger, Instant Clue: A Software Suite for Interactive Data Visualization and Analysis. Sci Rep 8, 12648 (2018).

90. S. Babicki et al., Heatmapper: web-enabled heat mapping for all. Nucleic Acids Res 44, W147–153 (2016).

91. H. Liu et al., Oncogenic HPV promotes the expression of the long noncoding RNA lnc-FANCI-2 through E7 and YY1. Proc Natl Acad Sci U S A 118, e2014195118 (2021).

